# A Morpho-Proteomic Atlas of Mitosis at Sub-Minute Resolution

**DOI:** 10.1101/2025.04.09.647158

**Authors:** Ede Migh, Vivien Miczán, Frederik Post, Krisztian Koos, Attila Beleon, David Kokai, Zsanett Zsófia Iván, Istvan Grexa, Nikita Moshkov, Reka Hollandi, David Csikos, Nora Hapek, Flóra Kaptás, Ferenc Kovacs, Andras Kriston, Diana Mahdessian, Ulrika Axelsson, Csaba Pal, Emma Lundberg, Mate Manczinger, Andreas Mund, Matthias Mann, Peter Horvath

## Abstract

Precise spatiotemporal protein organization is critical for fundamental biological processes including cell division^1,2^. Indeed, aberrant mitosis and mitotic factors are involved in diverse diseases, including various cancers^3,4^, Alzheimer’s disease^5^, and rare diseases^6^. During mitosis, complex spatial rearrangements and regulation ensure the accurate separation of replicated sister chromatids to produce genetically identical daughter cells^7–9^. Previous studies employed high-throughput methodologies to follow specific proteins during mitosis^10–15^. Still a temporally refined systems-level approach capable of monitoring morphological and proteomic changes throughout mitosis has been lacking. Here, we achieved unprecedented resolution by phenotypically decomposing mitosis into 40 subsections of a regression plane for proteomic analysis using deep learning and regression techniques. Our deep visual proteomics (DVP) workflow^16^, revealed rapid, dynamic proteomic changes throughout mitosis. We quantified 4,350 proteins with high confidence, demonstrating that 147 show significant dynamic abundance changes during mitotic progression. Clustering revealed coordinated patterns of protein regulation, while network analysis uncovered tight regulation of core cell cycle proteins and a link between cell cycle and cancer-linked mutations. Immunofluorescence validated abundance changes and linked previously uncharacterised proteins, like C19orf53, to mitosis. To facilitate data navigation, we developed Mito-Omix, a user-friendly online platform that integrates intricate morphological and molecular data. Our morphological and proteomic dataset spans mitosis at high resolution, providing a rich resource for understanding healthy and aberrant cell division.

## Main

Mitosis is associated with fundamental processes, including development, growth, and tissue repair, making it one of the most-studied phenomena in biology. A deeper understanding of the molecular mechanisms of mitosis promises to reveal the underlying causes of various diseases, particularly cancer, where mitotic dysregulation is a hallmark feature. Although a vast amount of data has been collected on the most important molecular players, this knowledge remains fragmented, and systems and network-level integration is lacking due to insufficient computational and experimental tools. Since bulk measurement of mitosis without chemical perturbations is challenging due to the natural asynchronicity between cells, and conventional immunohistochemical and microscopy methods only allow the concurrent analysis of 3-4 proteins, an adequate multiplexing solution is highly desirable. Stallaert et al.^17^ used iterative immunofluorescence of 48 pre-selected core cell cycle regulators to study cell cycle heterogeneity. Here, we present a detailed molecular and morphological dissection of human cell mitosis coupling 2D and 3D imaging with an objective artificial intelligence (AI)-based mitotic subsection recognition model with ultra-sensitive proteomic analysis. Previous works split the five major stages of mitosis into 8^18^ or 20 substages^13^. Another proteomic characterization of mitosis only acquired data on four out of five mitotic phases and focused more on the whole cell cycle^19^. We reasoned that further refinement of temporal and molecular resolution of mitotic changes would allow adequate modeling of protein dynamics and interaction networks. We therefore combined CAMI (Computer Aided Microscopy Isolation)^20^, DVP (Deep Visual Proteomics)^16^ and the Regression Plane concept^21^ with deep learning-based stage recognition to resolve mitosis into 40 specific consecutive subclusters that allow nearly continuous monitoring of protein quantities.

DVP allows the parallel measurement of up to 5000 unique proteins without pharmacological treatment, a significant advance over a previous analysis of 2000 proteins per FACS-sorted single cell measured using a cell cycle block and release experiment^22^. The single cell classification of the DVP workflow allowed us to accumulate cells with similar phenotypic features maximizing proteomic depth and accounting for heterogeneity between cells.

DVP analysis of batches of 120 single cells from the same subphase identified nearly 150 proteins that significantly changed in abundance throughout mitosis. Based on these changes, we clustered and functionally annotated proteins, followed by visualizing the connections between these proteins using a network-based approach. Our analysis also uncovered numerous proteins following similar molecular trajectories, implying they may have functional roles at specific mitotic stages. Overall, our approach provides a detailed and integrated morpho-proteomic map of mitosis, new molecular insights into mitotic regulation, and a flexible methodology that can be applied to other dynamic biological processes.

### AI-based, single-cell phenotyping and isolation for improved phenotypic decomposition of mitosis

Although mitosis is a continuous process, it is most frequently characterized by the five classical stages: prophase, prometaphase, metaphase, anaphase, and telophase. In order to observe the subtle mitotic events at the phenotypic level, we applied our Regression Plane (RP) concept^21^. We extended it using deep learning (DL)-based regression and classification techniques, in principle providing us with a computational tool to study continuous biological processes, including mitosis, at an arbitrarily fine resolution (Fig. 1a and Extended Data Fig. 1). We used HeLa cells stably expressing GFP-α-tubulin and H2B-mCherry to visualize structural changes in cellular morphology during mitotic progression. Time-lapse imaging of these cells was used to generate extensive temporal high-content imaging data, providing a precisely annotated training dataset for the RP. Single cells were segmented using NucleAIzer, our DL-based segmentation method^23^ and mitotic cells were annotated by experts who considered both morphological and temporal information to place cells at the appropriate position on the RP model of mitosis (Extended Data Fig. 1,2,3,4). This annotation strategy allowed us to represent the transitions of mitotic phenotypes in a continuous manner. An ensemble model of two deep convolutional networks was trained on the RP data and used for inference. Based on our measurements, the average prediction error was approximately 9 degrees around the 360-degree regression plane circle. Hence, we discretized the RP to 40 sub-stages of mitosis (Fig. 1a and Extended Data Fig. 1). Note that the 40 subsections are not isochronous but are based on the distribution of the five commonly accepted subphases according to our guideline set (shown in Extended Data Fig. 2). The complete mitotic event takes 93 minutes and, the quickest subsections last less than one minute (Fig. 1b, top) providing unparalleled resolution of mitotic morphology. As subsection 1 represents interphase, the distance in time from subsection 2 cannot be determined from the images due to the lack of marked morphological differences between individual interphase cells at different time points. To better understand the morphological changes over the course of the 40 subsections, in addition to the 2D widefield screens, 3D confocal image stacks were acquired and analyzed with the BIAS image analysis software, using a previously established automated workflow^24^ (Single-Cell Technologies Ltd.). This BIAS workflow enables automated correlative confocal screening by defining the position of each cell of interest using low-resolution wide-field images.

**Figure 1.**
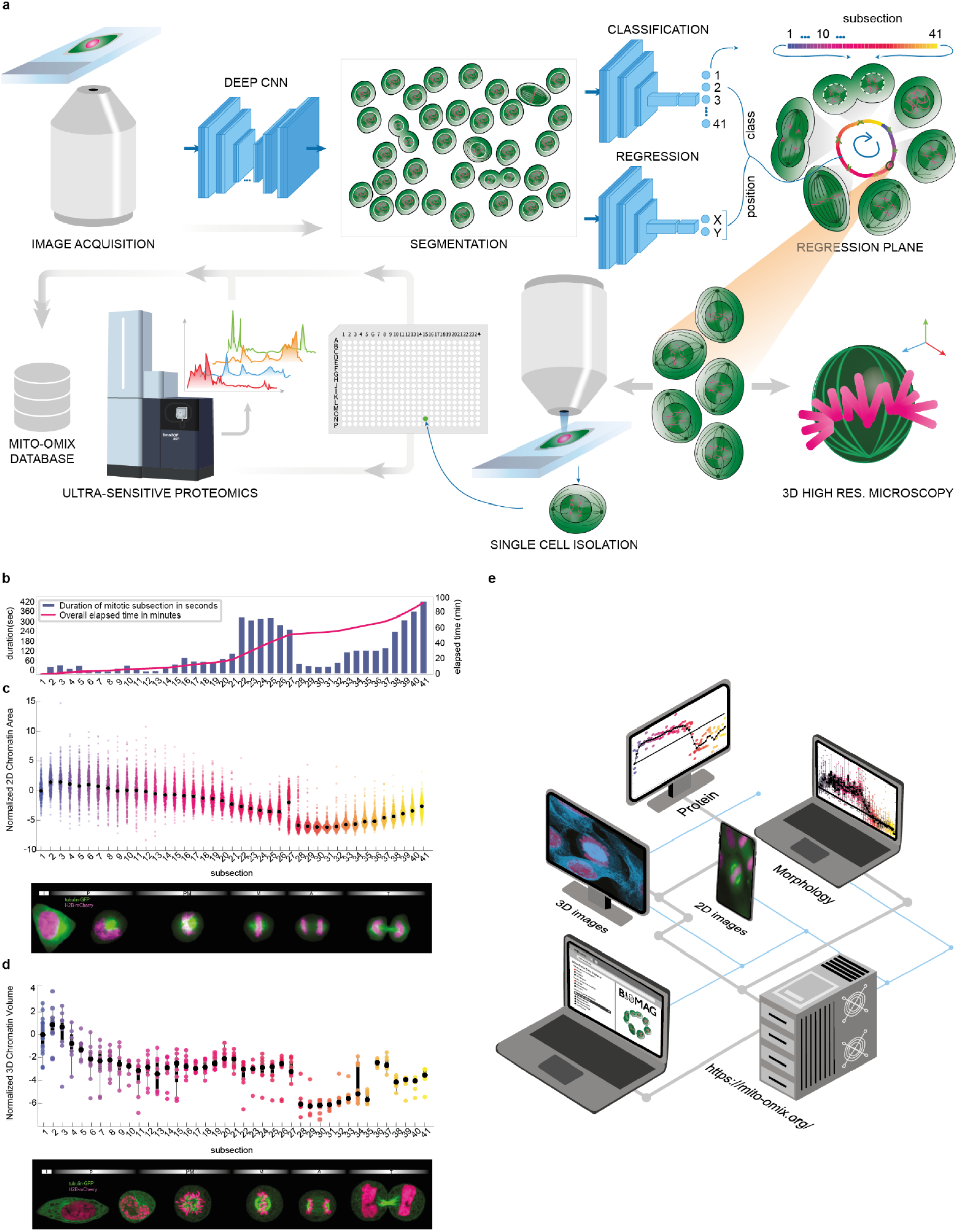
AI-based single-cell isolation workflow. a) Workflow consisting of high-throughput image acquisition, AI-based segmentation and classification/regression for the accurate labeling of 40 mitotic subsections, followed by 3D high-resolution confocal imaging and single-cell isolation for ultra-sensitive proteomic measurements. b) Temporal data on mitotic subsections. The blue bars represent the average duration of the individual mitotic subsections (in seconds, left axis). The overall elapsed time (in minutes, right axis) through mitotic subsections is represented by the red line. Example of the normalized 2D chromatin area feature (n=325-432 cells per group, groups 2,3 and 17-41 were significantly different from the interphase group, p<0.05, Kruskal-Wallis test with Bonferroni correction). c) Example of the normalized 3D chromatin volume feature. (n=9-20 cells per group, groups 11, 28-35 and 38-41 were significantly different from the interphase group, p<0.05, Kruskal-Wallis test). d) The MITO-OMIX database provides an interface to search, download and link the corresponding 2D/3D images, morphological information and proteomics measurements for the whole dataset.

Ultimately, using NucleAIzer to detect each cell and the RP to predict their mitotic stages (40 subsections+interphase) 120 cells per subsection were confidently predicted and then microdissected for proteomic measurements in 3 biological replicates (total of ∼41 x 120 x 3 = 14,760 selected and isolated single-cells). After analyzing the 2D and 3D segmented cells, gradual changes in features clearly indicated how the morphology of the cells progresses during mitosis (Fig. 1c, d). Each cell collected by microdissection is documented in the Mito-Omix database (Fig. 1d).

### Deep proteomic analysis of mitotic stages

To define the molecular landscape of the 40 mitotic classes, we used our recently introduced DVP approach for the proteomic analysis of cell populations. This approach allowed us to connect the single-cell image features and the regression plane model with mass spectrometry-based proteomics data based on cells selected by laser microdissection. The dissection of 120 cells per sample led to the quantification of 44,541 precursor peptides with a median of 30,221.5 precursor peptides and 5,735 protein groups across all samples and a median of 4,920 protein groups (Fig. 2a). 4,350 protein groups remained after a threshold of 67 % data completeness per replicate assuring accurate trajectories of protein intensities. The average median coefficient of variation of raw protein intensities across subsections was 15.7%, indicating that observed changes in protein levels are likely biologically meaningful rather than technical artifacts (Extended Data Fig. 5a). This consistency was also demonstrated by correlations of more than 0.8 or even 0.9 across samples (Extended Data Fig. 5b). Known markers of mitosis, BUB1, TOP2A, and TPX2^25–27^ and PCNA, a marker of DNA replication^28^, followed their expected trajectories along the 40 mitotic subsections, which highlights the accuracy of the experimental setup and validates the mitotic classifications (Extended Data Fig. 5c).

**Figure 2:**
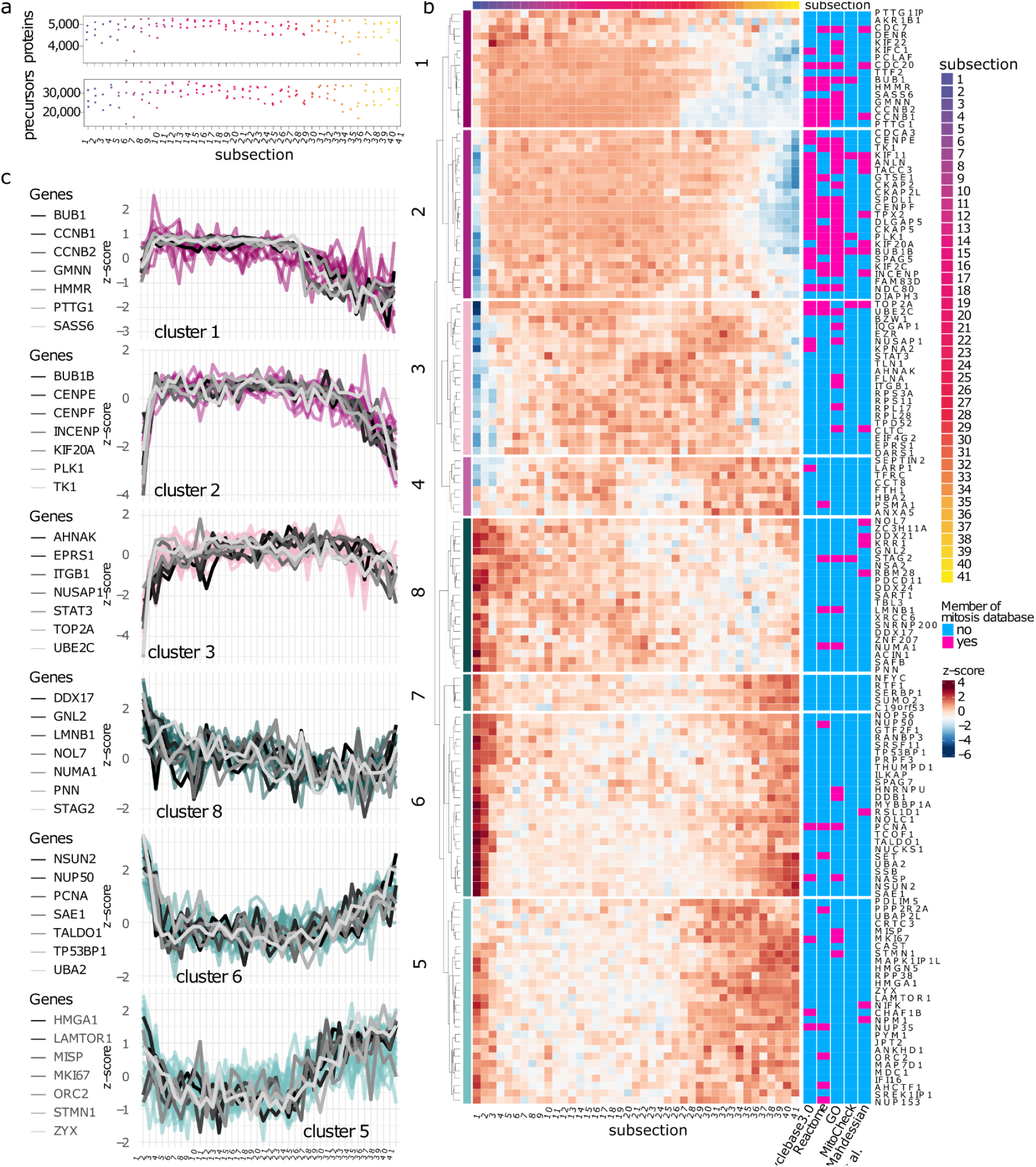
Proteomic landscape reveals protein variations across 40 mitotic classes. a) Quantified protein groups and precursors across 120 samples. b) Heatmap of significantly changed protein groups (Kruskal-Wallis FDR of 0.1). Proteins clustered into 8 distinct groups based on their z-score across the 40 mitotic subsections and the interphase. Row annotations demonstrate the presence of the protein group in the indicated commonly used mitotic resources. c) Abundance trajectories of clusters and highlighted proteins. Coloring of clusters corresponds to trends of up- (lines indicated by pink color) and down-(lines indicated by cyan color) regulated clusters of proteins.

### Analysis of fine-grained mitotic stages reveals 147 significantly changing proteins and major tendencies of change

The acquisition of proteomes of 40 mitotic classes allowed us to characterize protein dynamics at an exceptionally high temporal resolution of less than 1 minute and an average resolution of less than 2 minutes (Fig. 1b top). 147 proteins were identified that significantly changed their abundance between the 40 mitotic subsections and interphase (Fig. 2b, Extended Data Table 1). The increased temporal refinement of proteomic changes allowed a number of insights into mitotic mechanisms. Approximately half of these significantly changing proteins increased their abundance during the first subsections of mitosis compared to interphase (Fig. 2c, clusters 1-4). The overlap with other mitotic resources^13,14,29–31^ revealed that proteins with an early increase in abundance tend to be associated with known mitotic functions (Fig. 2b right). Of note is that the observed increases in abundance in clusters 2 and 3 were relatively abrupt while decreases in later mitotic phases were more gradual. The three kinetochore proteins CENPE, CENPF, and INCENP, known to be essential for chromosome segregation and cytokinesis were located in this cluster^32,33^. Despite having similar functions, CENPE decreased in subsection 36, CENPF in subsection 37, and INCENP in subsection 39. It was previously described that kinetochore subunits are dephosphorylated leading to disassembly^34^. Future research should investigate if the observed consecutive protein degradation indicates that the disassembly follows an order in which CENPE is prioritized over CENPF and INCENP.

A particularly large decrease in abundance occurred for proteins in cluster 1 between classes 26 and 27. These proteins include the DNA replication inhibitor Geminin (GMNN) which is known to be ubiquitinated by the anaphase-promoting complex (APC) allowing DNA replication during S phase^35^. Protein degradation by the APC appeared to take only a few minutes and was highly specific for GMNN, PTTG1, CCNB1, and CCNB2. Other known targets of the APC like CDC20 or BUB1B did not decrease in the same manner. Known interaction partners, like PTTG1 and PTTG1IP, also showed slightly different changes in abundance pattern. PTTG1IP decreased at subsection 33, well after PTTG1 levels declined at subsection 27. Upon interaction of PTTG1IP and PTTG1, the complex is known to be associated with chromatin enabling the interaction of PTTG1 with the DNA^36^. This interaction is essential for its function as an inhibitor of ESPL1. By cleaving cohesin, ESPL1 enables the separation of sister chromatids during anaphase^37^.

The previous paragraph focussed on proteins that are essential for cell division, which are expected to be upregulated during the early phases of mitosis. On the other hand, proteins that decrease in abundance are likely to be either inhibitors of cell division or proteins that lose their function during mitosis, e.g., DNA replication proteins. 79 proteins displayed a decrease of abundance during the first phases of mitosis (cluster 5-8). About a third of these proteins were previously recorded in other mitotic resources. The cohesin subunit STAG2 is important for kinetochore-microtubule attachment and localization of BUB1^38^. STAG2 increased in abundance from subsection 3 to 7 and steadily decreased until subsection 20. BUB1 and other proteins that are related to the kinetochore are located in cluster 1 and only decreased after subsection 27. STAG2 is known to be degraded by prophase pathways, suggesting that it is especially crucial for the cohesin complex in facilitating the attachment of microtubules to kinetochores^39^. Cluster 5 consisted of proteins that were decreased at the beginning of mitosis but increased again at subsection 27, opposing the tendencies in cluster 1, which consisted mostly of important mitotic proteins. Part of this cluster were the nuclear pore complex proteins NPM35 and NPM153, which shows that the translation of these proteins is starting before the assembly of the nucleus at the end of the telophase.

Protein abundance fluctuated throughout mitosis if looked subsection-by-subsection however, the global trend of protein abundance in clusters 5, 6 and 8 went down during mitosis from initially high abundance in interphase and then restored back in telophase. Abundance of proteins in clusters 2 and 4 had the opposite trend. Proteins of cluster 1 had a similar trend to 2 and 4, though the measured protein abundance at the end of mitosis was lower than in interphase. That might indicate that the abundance of cluster 1 proteins was restored during another phase of the cell cycle. Some of the proteins in cluster 8, i.e., SAFB, XRCC6, and DDX17, were related to the DNA damage response. SAFB transiently binds to damaged chromatin to mediate DNA damage signaling^40^. XRCC6 performs non-homologous DNA end-joining to repair DNA double-strand breaks ^41^. Even though DDX17 is mainly known for RNA processing, it was also shown to be involved in the DNA damage response through interaction with HDAC1^42,43^. Other DNA damage response proteins such as PCNA or TP53BP1 were located in cluster 6 and were only increased in the first subsections of mitosis and towards the end of mitosis^44,45^.

### Grouping of subsections identifies more than 1000 significantly changed proteins

Our initial analysis involved 41 time points with 3 replicates at each time point. This approach allowed us to capture fine-grained changes in protein abundance during mitosis. By aggregating time points, we increased the statistical power of the data resulting in the identification of 1060 significantly changed proteins across 14 subsections (Extended Data Fig. 6a). These proteins were grouped into 10 clusters with distinct biological processes. While mitotic proteins were increased from subsection 5 to 35 (cluster 8), DNA replication (cluster 2) and RNA processing (cluster 3) were decreased during these subsections. Histone-related proteins (cluster 6) were increased from subsections 21 to 38. Nuclear pore complex proteins increased steadily from subsection 27 to 41, before the nucleus is assembled or newly synthesized histones are needed for DNA replication^46^. These increases might seem ahead of time, but z-scores can sometimes amplify differences, indicating a stronger change than what is evident from the actual intensities, due to their reliance on standardized deviations relative to the dataset’s variability. A selection of proteins linked to these different biological pathways were plotted separately to highlight their mitotic dynamics (Extended Data Fig. 6b-j). The mitotic kinases AURKA, AURKB, BUB1B, CDK1, and PLK1 increased between interphase and the mitotic subsections 2-4 (Extended Data Fig. 6d). BUB1 was found to have a higher relative abundance in interphase than the other mitotic kinases. Furthermore, BUB1 decreased already at subsection 30, which was the earliest of all mitotic kinases. CDK1 remained on high levels even at the end of mitosis. Different abundances during interphase were also observed for centromeric proteins (Extended Data Fig. 6f). While CENPC and CENPV decreased at the beginning of mitosis, CENPF, CENPE, and INCENP increased. Cytoskeletal proteins like ACTN1, ACTN4, and TUBB6 increased during mitosis (Extended Data Fig. 6h). In contrast, the nuclear skeletal proteins LMNA, LMNB1, and LMNB2 decreased during mitosis, which corresponds to the degradation of the nuclear envelope seen by microscopy (Fig. 1b-c), and started reestablishing from subsections 39-41 (Extended Data Fig. 6h). The proteasomal subunits PSMA1, PSMA3, PSMC1, PSMC6, and PSMD14 showed an overall increase in abundance during mitosis (Extended Data Fig. 6b). However, this increase was interrupted by a decrease between subsection 21 and subsection 26. These subsections are representing the metaphase in which mitosis remains for almost 30 minutes before chromosomes are separated (Fig. 1b). Several ribosomal subunits strongly increased in abundance between subsection 8-10 and subsection 11-13 (Extended Fig. 6g). A different regulation was observed for mitochondrial ribosomal subunits that decreased from interphase to the start of mitosis but proceeded with a steady increase towards the end of mitosis.

### The functional association network of proteins with changing abundance throughout mitosis

To further analyze the functions of the proteins with significantly changing abundance during mitosis, we turned to the STRING database^47^ and the Cytoscape software^48^. We aimed to uncover whether proteins that exhibit similar expression patterns during mitosis also cluster together in a network characterized by functional associations and physical interactions (Fig. 3a). This additional clustering resulted in proteins belonging to two major and multiple minor clusters. Proteins enriched in the first major cluster were associated with the mitotic cell cycle process, while those in the second dominantly took part in rRNA metabolic processes. Importantly, proteins belonging to the functional clusters showed similar expression patterns. For example, 41 of 52 proteins (79%, Odds Ratio (OR): 6.58 vs. proteins outside cluster, Fisher’s exact test P: 5.37×10^-7^) from the first functional cluster belonged to expression clusters 1 to 4 with increasing abundance at the beginning of the mitosis, while 20 of 21 proteins (95%, OR: 26.04 vs. proteins outside cluster, Fisher’s exact test P: 5.995×10^-6^) from the second functional cluster were found in expression clusters 5 to 8.

**Figure 3.**
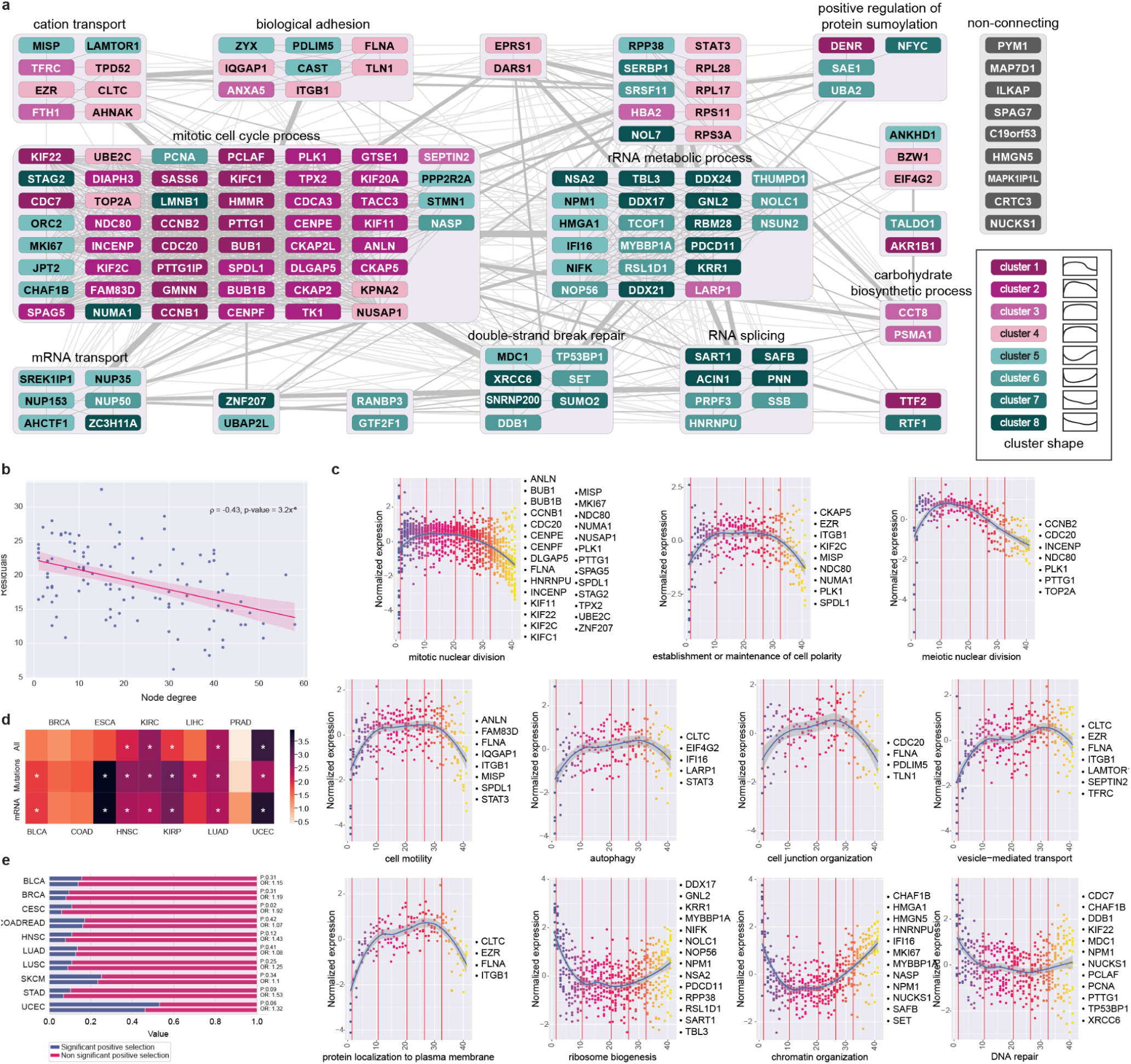
Functional analysis of significantly changing proteins. a) Protein-protein interaction network of the 147 proteins that change significantly during mitosis. Interactions include direct (physical) and indirect (functional) associations; Nodes in different interaction clusters are in separate boxes. Clusters are named after the corresponding enriched GO terms. Node colors represent expression clusters on Fig. 2c. Edge thickness indicates the interaction density between different clusters. b) The sum of residuals, used as a measure of the strictness of expression regulation during mitosis, is plotted against the node degree. The red line indicates a linear regression line with 95% confidence interval. Spearman’s rho and two-sided P value of the correlation test is shown. c) The scatterplot shows the expression of proteins belonging to different GO process groups in different stages of mitosis. The blue curve indicates a regression line fitted using 3rd degree polynomials and the gray area indicates 95% confidence interval. d) The heatmap displays the odds ratios (ORs) for the likelihood of identifying driver genes among mitosis-associated versus non-mitosis-associated genes in different tumor types. Asterisk (*) indicates statistical significance with a one-sided P value less than 0.05 according to Fisher’s exact tests. e) The plot presents the proportion of positively selected genes among mitosis-associated and non-mitosis-associated genes across various tumor types. The ORs and the one-sided P values of Fisher’s exact texts are indicated. BLCA: Bladder Urothelial Carcinoma, BRCA: Breast Invasive Carcinoma, CESC: Cervical Squamous Cell Carcinoma and Endocervical Adenocarcinoma, COAD: Colon Adenocarcinoma, ESCA: Esophageal carcinoma, HNSC: Head and Neck Squamous Cell Carcinoma, KIRC: Kidney renal clear cell carcinoma, KIRP: Kidney renal papillary cell carcinoma, LIHC: Liver hepatocellular carcinoma, LUAD: Lung Adenocarcinoma, LUSC: Lung Squamous Cell Carcinoma, PRAD: Prostate adenocarcinoma, READ: Rectum Adenocarcinoma, SKCM: Skin Cutaneous Melanoma, STAD: Stomach Adenocarcinoma, UCEC: Uterine Corpus Endometrial Carcinoma;

Prior research underscores that key regulators of mitosis are subject to stringent control of gene expression^49,50^. Our extensive dataset on protein abundance enables us to analyze the dynamics of protein expression along mitosis. Based on this, we hypothesized that central proteins in our network would display more tightly regulated expression patterns. To evaluate this, we applied polynomial regression - an established method for analyzing gene expression in time-course experiments^51^ and fitted regression models to the scaled data as a function of pseudo-time to infer the dynamics of protein abundance during mitosis. We then calculated the sum of the absolute values of the residuals for each protein (see Methods). Lower sums of residuals indicate more consistent protein levels and tighter expression regulation, whereas higher sums point to more stochastic variations, implying less stringent regulation. Our findings confirm that central proteins in the functional network demonstrate more strictly regulated expression changes throughout mitosis (Fig. 3b). The results remained significant when we controlled for absolute protein expression and biological replicates in a multivariate linear regression model (Extended Data Table 2). As expected, proteins within the “mitotic cell cycle process” cluster exhibited the highest degree centrality and displayed the lowest residual sums, followed by the “rRNA metabolic process” cluster. Accordingly, proteins belonging to these clusters showed the lowest residual sums (Extended Data Fig. 7).

To identify protein functions associated with specific expression patterns, we examined proteins belonging to different functional GO terms (Fig. 3c). We saw two major trends in protein levels according to functional groups. The abundance of proteins in the first main group initially increased and then declined. Within this group, some functional clusters had smoother expression patterns: mitotic nuclear division, chromosome segregation, establishment and maintenance of cell polarity and cell motility, whereas more biphasic and skewed trajectories were found for clusters: autophagy, cell junction organization, vesicle-mediated transport, protein localization to plasma membrane and meiotic nuclear division. In contrast, three GO groups, namely ribosome biogenesis, DNA repair and chromatin organization showed an opposite pattern. This suggests that proteins with these functions are downregulated during mitosis.

### Proteins engaged in mitotic process and their relation to cancer-related pathways

Dysregulation of the cell cycle is a fundamental aspect of cancer development^4^. Studies indicate that mitosis-associated genes often exhibit altered expression in tumors more frequently than they harbor specific driver mutations^3,52^. We analyzed driver gene data of DriverDBv4, which aggregates not only mutational information but also data on expression outliers and other omics layers^53^. For each cancer type, we selected all identified driver genes in the database and matched them with the ones encoding proteins in our dataset. We then assessed the prevalence of driver genes in two distinct groups: those encoding proteins associated with mitosis and those that are not. Mitosis-associated genes were more frequently identified as driver genes in 5 out of 11 tumor types (Fig. 3d, Fisher’s meta P: 1.07×10^-9^). This trend became even stronger when the analysis was carried out separately on two subsets: one for driver genes with specific mutations (Fig. 3d, significant enrichment in 8 out of 11 tumor types, Fisher’s meta P: 1.42×10^-9^) and another for those exhibiting altered gene expression in tumors (Fig. 3d, significant enrichment in 6 out of 11 tumor types, Fisher’s meta P: 2.01×10^-13^ heatmap values are listed in Extended Data Table 3). Importantly, in the majority of tumor types where the results were not statistically significant, the observed trends still aligned with those seen in other cancer types, suggesting a consistent pattern. The only exception was prostate adenocarcinoma, which may indicate distinct mechanisms of oncogenesis that predominate in this tumor type. We also investigated whether driver genes are overrepresented within any of the previously analyzed functional clusters compared to others. After adjusting for multiple hypothesis testing, we found no significant associations, indicating that driver genes are likely to be evenly distributed across the various functional clusters of mitosis-associated genes (Extended Data Table 4).^52^

Our previous findings remained consistent when we applied the Cancer Bayesian Selection Estimation (CBaSE) tool^54^ to assess the selection pressure on genes across various cancer types within The Cancer Genome Atlas (TCGA) database (see Methods for details). Notably, there was a significant enrichment of positively selected genes within the mitosis-associated group in cervical squamous cell carcinoma (OR: 1.92, P value of one-sided Fisher’s exact test: 0.02), and we observed similar, though non-significant, trends in other cancer types (Fig. 3e, Fisher’s meta P: 0.015).

### Fluorescent antibody imaging to complement proteomics analysis during mitosis

To enrich the mass spectrometry-based measurements with localization information, we combined RP-guided, mitotic phenotyping with immunofluorescence staining for proteins of specific interest.

First, we selected cyclin B1 (CCNB1), a well-known protein required for mitotic processes such as chromosome condensation, nuclear envelope breakdown and spindle pole assembly. Consistent with protein abundance measurements and former publications^55^, the fluorescent signal intensity for CCNB1 rapidly increases during the first mitotic subsections, while in the 27th mitotic subsection, when sister chromatid separation initiates, staining suddenly disappears or significantly decreases (Fig. 4a). These results independently validate that our workflow can capture one-minute snapshots of changes during mitosis based on a proteomic profile. Subsequently, to corroborate the mitotic hit list, we examined uncharacterized proteins associated with mitosis. Our first candidate, EPRS1, is a Glutamyl-prolyl-tRNA synthetase which charges tRNAs with their cognate amino acids. A previous publication^56^ showed that knockdown of EPRS1 in hepatocellular carcinoma (HCC) cells such as Huh7 and Hep3B dramatically suppressed HCC cell proliferation rate, potentially implicating EPRS1 in cell division. Consistent with this hypothesis, both protein abundance and fluorescence intensity measurements showed elevated EPRS1 gene expression level during mitotic phases suggesting it may function during mitosis (Fig. 4b).

**Figure 4.**
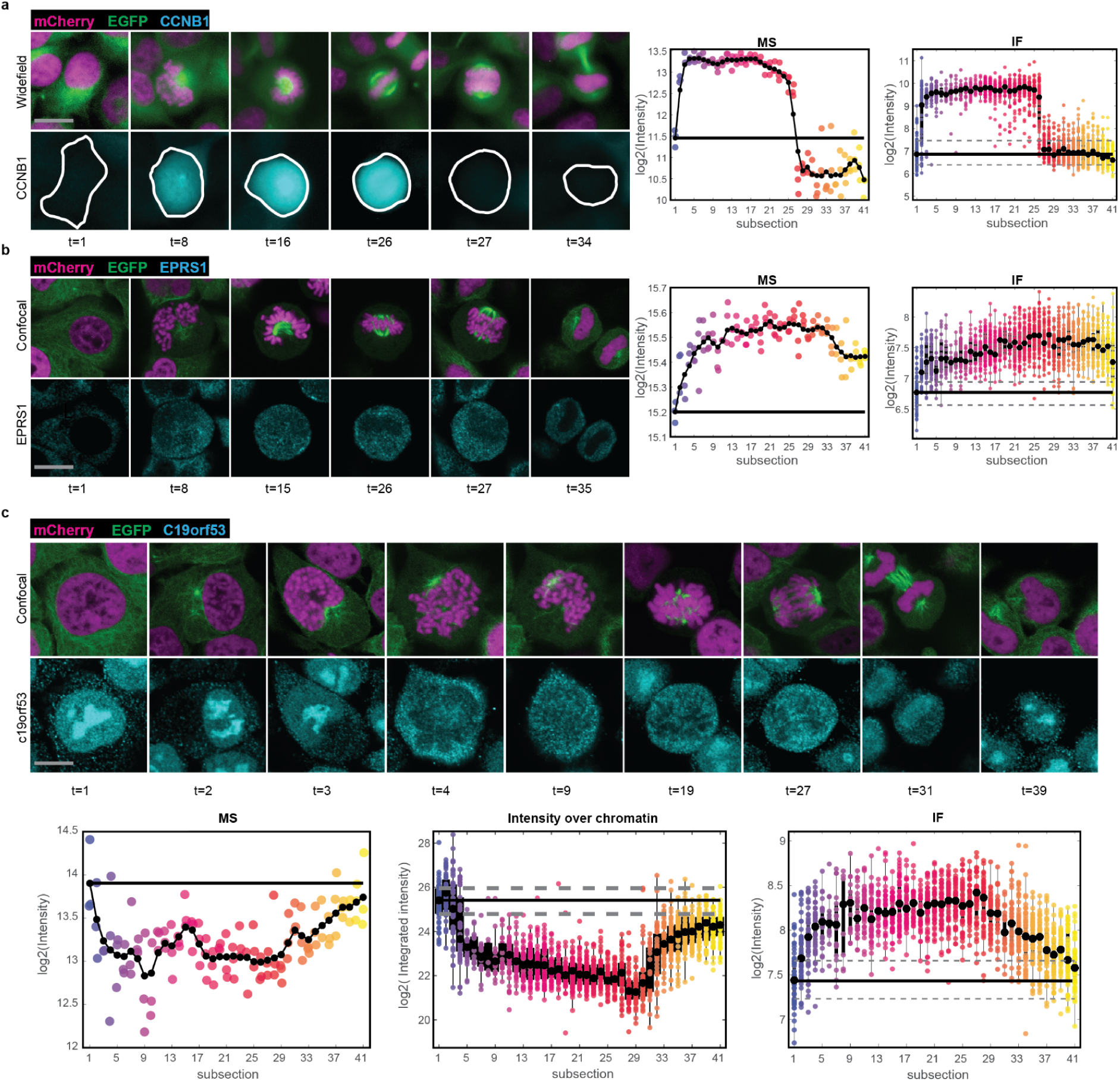
Complementation of proteomic measurements with immunofluorescence staining. a) Left: Widefield microscopy images of CCNB1 (cyan) immunostaining on HeLa Kyoto cells expressing EGFP-alpha-tubulin (green) and H2B-mCherry (magenta) at different mitotic subsections. Middle: Proteomic measurement of CCNB1 protein over interphase plus 40 mitotic subsections in three biological replicates. Right: average CCNB1 immunofluorescence intensity. b) Left: Confocal microscopy images of EPRS1 (cyan) immunostaining on HeLa Kyoto cells expressing EGFP-alpha-tubulin (green) and H2B-mCherry (magenta) at different mitotic subsections. Middle: Proteomic measurement of EPRS1 protein during interphase plus 40 mitotic subsections in three biological replicates. Right: average EPRS1 immunofluorescence intensity. c) Top: Confocal microscopy images of C19orf53 (cyan) immunostaining on HeLa Kyoto cells expressing H2B-mCherry (magenta) at different mitotic subsections (EGFP channel not shown). Lower left: Proteomic measurement of C19orf53 protein over interphase plus 40 mitotic subsections in three biological replicates. Lower middle: average C19orf53 immunofluorescence intensity over the chromatin area. Lower right: average C19orf53 immunofluorescence intensity in the whole cell. Scale bars: 10 µm in (a) and (b); 5 µm in (c).

Next, we assessed the abundance of the Leydig cell tumor 10 kDa protein homolog (C19orf53), a factor previously linked to ribosomal assembly^57^ and suggested to mediate cell proliferation^58^. High-resolution 3D confocal images revealed that in interphase this protein is localized in the nucleus (most likely in the nucleoli). As mitosis progresses, the quantity of C19orf53 decreases, and it redistributes to the cytoplasm, reverting to its original localization state during the late phases of mitosis (Fig 4c). Interestingly, protein abundance measurements show downregulated expression during mitotic phases which correlates with the intensity measurement over the chromatin channel (Fig. 4c).

Overall, we performed the immunostaining experiment for 15 proteins, the 3 described above (C19orf53 *r_s_* =− 0. 651, *P* < 0. 001; CCNB1 *r_s_* = 0. 720, *P* < 0. 001; EPRS1 *r_s_* = 0. 460, *P* < 0. 001) and 12 additional proteins. For 6 out of 12 of additional proteins we detected the expected correlation between antibody staining intensity and protein abundance (INCENP *r_s_* = 0. 599, *P* < 0. 001; MAPK1IP1L *r_s_* =− 0. 476, *P* < 0. 001; PTTG1 *r_s_* = 0. 754, *P* < 0. 001; SUMO2 *r_s_* =− 0. 522, *P* < 0. 001; TLN1 *r_s_* = 0. 782, *P* < 0. 001; UBE2C *r_s_* = 0. 447, *P* < 0. 001; ). In 6 out of 12 proteins intensity and measured protein abundance did not correlate, which could be due to antibodies detecting only a subset of protein isoforms and 3D structures, or antibodies not penetrating the cell compartment of protein localization (Extended Data Fig. 8, 9).

### MITO-OMIX: a comprehensive repository of mitotic data

We developed a data portal to provide quick and easy access to our measurements (https://mito-omix.org/, Fig. 1d). MITO-OMIX contains 2D and 3D morphology features, including access to every image of an isolated single cell as well as 3D data stacks. These data are integrated with access to the proteomics measurements. Data can be queried by feature names in the case of the morphology data and by gene names of the proteins. It is possible to query the immunofluorescent staining data and images. MITO-OMIX also includes links to useful databases (e.g. HUGO Gene Nomenclature Committee (HGNC), Ensembl, Uniprot, Gene Ontology (GO) and Human Protein Atlas (HPA)).

## Discussion

In this study, we report a cutting-edge, high-throughput AI-driven workflow designed for the comprehensive analysis of mitotic cell dynamics that seamlessly integrates morphological changes and molecular contributors with unparalleled precision and resolution. Importantly, this image analysis-based methodology is versatile and promises to extend the exploration of various dynamic cellular processes (eg. mitochondrial dynamics, Golgi vesicle transport, apoptosis or autophagy) in a more natural and physiologically relevant context. Unlike previous studies that relied on cell sorting or chemical treatments to accumulate cells with certain phenotypes, we did not artificially arrest the cell cycle to obtain mitotic cells, allowing for the observation of cells in their native state and avoiding the induction of artifacts associated with such treatments or the sorting procedure ^19,59,60^. The approach has yielded a unique proteomic resource of mitotic processes: the abundance of 147 proteins changed significantly between the 41 subsections, revealing intricate regulation patterns like the degradation of mitotic regulators as CCNB1 that decreased 5-fold in abundance within the first minute of anaphase. The temporal resolution of our data allowed us to identify putative priorities in this APC-dependent protein degradation^61^. Other APC targets like AURKA, AURKB, BUB1B, CDC20, PLK1, TK1, or TPX2 were not degraded within the first minute of anaphase but more gradually (Fig. 2b,c; Extended Data Fig. 6d,e)^62–66^. The targeting of the APC is dependent on the APC-subunits CDC20 and FZR1^67^. CDC20 was suggested to be required for metaphase-anaphase transition, which is in line with its decrease starting between the subsections 31 and 34 that correspond to the end of the anaphase (Fig. 2b). The degradation of proteins during the exit of mitosis was suggested to be dependent on FZR1. Despite not being significantly changed in abundance in our data, FZR1 increased in abundance towards the end of anaphase and telophase. The high resolution of our regression plane model in combination with the DVP technology allowed us to observe these transient processes, which were discovered through numerous meticulous studies, within a single experiment. We thereby provide a complementary perspective to previous research on mRNA regulation and translation during the cell cycle. While it has been shown that mRNA transcription and translation is generally repressed during mitosis, certain mRNAs maintain higher translational activity^59^. It is broadly understood that increased protein abundance is more likely to correlate with high mRNA levels and elevated translation rates, whereas low protein abundance can arise from multiple factors beyond decreased translation, such as protein degradation or reduced stability^68^. A second study on mRNA abundance during mitosis and transition to G1 observed regulation that were mainly associated with mitotic exit, cell-type specific genes, and DNA replication, not capturing mitotic drivers^60^.

By regrouping the mitotic subsections from 41 to 14 time points, we increased the statistical power of our analysis, which yielded in 1060 proteins with potential roles in mitosis. Among these proteins, we found different abundance patterns of the centromere subunits CENPC, CENPE, CENPF, CENPV, and INCENP (Extended Data Fig. 6f). CENPC and CENPV were highly abundant during interphase and decreased in the first subsections of mitosis. Both of these proteins are essential for the correct assembly of the centromere: CENPC functions as a hub for centromere assembly at the kinetochore^69,70^, and CENPV depletion leads to mislocalization of the centromere^71^. Furthermore, proteasomal subunits increased during mitosis with the exception of a decrease between subsections 21 to 26, which corresponds to metaphase (Extended Data Fig. 6b). Inhibition of the proteasome was shown to lead to an aggregation of centromeric proteins in the center of the cell, putatively due to failure of degrading polyubiquitinated proteins^72^. The aggregates dissolve after removal of the proteasome inhibitor. The same mechanism of attenuated proteasomal activity could assure that chromosomes are correctly attached to kinetochores during metaphase before transitioning to anaphase.

Thorough examination of the functions and interconnections within the selected protein cohort further strengthened our understanding of their involvement in the process. Moreover, our findings suggest that the genes encoding these proteins are likely to be key drivers across multiple cancer types. Notably, the most pronounced enrichment was observed in esophageal cancer, corroborating previous research that highlighted the critical role of the gene CCNB1 in this disease^73^. Additionally, we identified significant enrichment in two HPV-associated cancers—cervical squamous cell carcinoma and head and neck squamous cell carcinoma—as well as in kidney cancers (renal cell and papillary cell carcinoma), uterine corpus endometrial carcinoma, lung adenocarcinoma, liver hepatocellular carcinoma and bladder cancer. These results emphasize the clinical significance of our study and the potential therapeutic targets within these protein clusters.

Although a significant portion of the regulation during mitosis is attributed to post-translational modifications that do not necessarily result in alterations at the protein expression level^74^, several hits of proteins with significantly changing abundance were validated by immunofluorescence staining coupled with high-resolution 3D confocal microscopy, showing that our approach is capable of retrieving uncharacterized players in mitosis. As one example, the uncharacterized protein C19orf53 was highlighted by our approach, suggesting its likely role in mitosis. We observed that C19orf53 was colocalizing with the nucleolus and the validation of MS quantification only held true for this colocalizing variant of C19orf53. This indicated a structural change or post-translational modification of C19orf53 between subsection 4 and 27 that possibly obscured the epitope. Ubitiquinations and phosphorylations of the targeted epitope of C19orf53 have previously been reported^40,75–77^. Further experiments are necessary to clarify the localization of C19orf53 and putative structural changes during mitosis.

In conclusion, this workflow provides a map of mitosis with high temporal resolution. Along with our follow up analyses, which present broadly applicable approaches to integrating and analyzing large datasets, our results have unlocked new avenues for the in-depth analysis of mitotic events within the complex context of human tissues. These data may ultimately help identify and understand differences in cell division based on tumor-type and even at the patient-specific level.

The major novelty of our approach lies in the deep learning-based identification of an unprecedented number of 40 mitotic subsections combined with the exceptional proteomic depth. This enabled sensitive quantification of proteins essential for cell survival or with redundant functions, whose mitotic significance cannot be studied in a trivial manner by single gene inactivation studies. The information gathered here on the molecular, morphological and physical properties of mitosis is a community resource that we share in a public database with the aim to help understand the role of mitosis in human diseases such as cancer. Even novel targets for treatment of those diseases can be uncovered, for instance, cell division regulator drugs already exist and are used in cancer therapy but whose exact mechanisms of action are unknown ^78,79^. The approach we present is extendable to human tissue slices and gives the opportunity to study specific cancer types even on the patient-level .

## Methods

### In vitro experiments

The human endocervical adenocarcinoma HeLa Kyoto EGFP-alpha-tubulin/H2B-mCherry (CLS GMBH, 300670) cell line was grown in modified Dulbecco’s modified Eagle’s medium (DMEM) containing high glucose (4,5 g/l), 2mM L-glutamine, 10% Fetal Bovine Serum (Gibco), 0.5 mg/mL G418, and 0.5 mg/mL puromycin at 37°C in a 5% CO_2_ humidified environment. These cells are endogenously tagged with two fluorescent proteins fused to the microtubule protein alpha-tubulin and the H2B histone to visualize the morphological changes in the cytoskeleton and the chromatin, respectively.

In order to enrich for HeLa cells undergoing mitosis, a mitotic shake-off protocol was applied. Namely, cells were grown in a flask to a subconfluent phase, then, detached mitotic cells were removed by gentle shaking of the flask in a new medium and transferred to a sterile tube. After brief centrifugation (5 mins 1510 rpm at 24°C) the supernatant was removed and the cells were resuspended in pre-warmed DMEM and cultured on PET frame slides (Leica, 11505151) pretreated with EtOH and UV irradiation for 30 minutes. After cultivating cells for 24 hours on PET frame slide, cells were fixed for microscopy experiments.

Cells prepared for the proteomics experiments were fixed for 10 minutes in 4% paraformaldehyde (PFA), then thorough washing was performed in Phosphate-Buffered Saline (PBS) and purified water. In contrast, cells prepared for the training set for the regression plane were fixed promptly after the live-imaging session with the same method.

### Immunofluorescence staining for localization experiment

Targets for immunofluorescence staining were selected based on literature data and antibody availability in the Human Protein Atlas (HPA) project. Proteins whose abundance during mitosis is known were selected as “positive controls”, while uncharacterized proteins lacking literature or an association with mitosis were chosen as “interesting hits”. The fixation and staining procedure were performed as previously described^80^. Briefly, after screening cells were fixed for 10 minutes in 4% PFA. Fixed cells were then washed twice in PBS and permeabilized using 0.25% Triton X-100 in Tris-buffered saline (TBS) (0.05 M Tris-HCI, 0.15M NaCl, pH 7.6). Blocking of nonspecific binding sites was performed in PBS containing 5% Bovine serum albumin for 1 hour. Primary antibodies were diluted in PBS containing 1% BSA (PBS-B) and incubated overnight at 4°C. HPA IDs of primary antibodies are listed in Extended Data Table 5. For secondary antibody staining, goat anti-rabbit secondary conjugated Alexa 647 antibody (Thermo Fisher, 711-605-152) was diluted in PBS-B (1:600) and incubated for 1 hour at room temperature.

### High-content screening

Cells were then imaged using a PerkinElmer Operetta High Content Screening system controlled by Harmony 3.5.1 software. In the case of the training images two-channel time-lapse image stacks of two focal planes were acquired with 20× 40× and 60× objectives. For each field of view, a 100-frame image sequence was acquired with a 2-minute gap between subsequent frames for the regression plane dataset and only one time-point was used for the omics experiments using a 60× objective. The system was maintained at a constant temperature (37 °C) and CO_2_ concentration (5%) for optimal cell growth. The following filter set was used: for EGFP excitation was 460-490 nm and emission was 500-550 nm, for mCherry excitation was 520-550 nm and emission was 580-650 nm and for immunofluorescence staining (Alexa 647) excitation was 620-640 and emission 650-700. Excitation was provided by a Xenon lamp. Live-cell imaging was performed in DMEM, while PBS was used as an imaging medium in the case of fixed samples.

### Image processing

Raw high-content screening images were processed using the Biology Image Analysis Software (BIAS)^24^. Since the regular interphase cells and the slightly rounded-up mitotic cells appeared in focus on different planes, we first generated a maximal intensity projection of the two different focal planes. Then, illumination correction was performed with the built-in CIDRE^81^ method. Segmentation of the nucleus and cytoplasm channels was achieved with built-in deep neural network models (cytoplasm: Generic Cytoplasm segmentation v1, input spatial scaling parameters for 20×, 40× and 60× images 1.0, 0.7, 0.4, respectively; nucleus: Generic Nucleus Segmentation v1, input spatial scaling parameters for 20×, 40×, 60× images 2.4, 2.4, 1.0, respectively), and minimal object size constraints were used (500 and 1000 pixels, respectively). The resulting object masks touching the image borders were filtered out, then cytoplasm masks were dilated by 1 um to minimize loss of cell material due to laser microdissection. Cytoplasm and nucleus masks were matched and image features (see MITO-OMIX database for the visualization of feature data) were extracted from them. In cases when we used statistical tests to evaluate differences between mitotic stages, the Kruskal-Wallis test was used with Bonferroni correction to compare groups, as according to Shapiro-Wilk tests, not all groups were normally distributed, so nonparametric tests were used for comparison of groups.

Image features were fed to a pre-trained machine learning classification model to filter out image artifacts. Cropped images of 150×150 pixels were then generated from the remaining objects for deep learning purposes.

### Confocal imaging

10 cells from each mitotic subsection were randomly selected for high-resolution 3D confocal imaging using a Leica SP8 setup controlled by LasX software. Registration of the fields of view of the Operetta and SP8 was performed using marker coordinates engraved in the PET membrane to allow easier detection of the selected cells. 512×512 two-channel image stacks were acquired using 6-15 focal planes with 0.09×0.09×2 um voxel size utilizing a high-NA(1.2) water-immersion 63× objective. EGFP was excited at 588 nm and detected at 495-550 nm, mCherry was excited at 552 nm and detected at 650-775 nm, Alexa 647 was excited at 638 nm and detected at 650-695 nm wavelengths in sequential mode to avoid crosstalk.

### Deep learning model for cell selection

An image database was generated for training of the neural networks. 40×, and 60× objectives were used for acquiring time-lapse images of 309 mitotic cells (131, and 178 cells at each magnification). This resulted in 11797 image crops (6868, and 4929, respectively).

Annotation of the image sequences was performed using the Advanced Cell Classifier (ver. 3.1) software. Cells were identified by a standard tracking algorithm (simpletracker^82^, with alignment from CellTracker^83^) on consecutive image frames and cropped cell images were placed on a circle with a 4500 pixel radius lying on a square of 10000×10000 pixels according to a previously defined annotation strategy (Extended Data Figure 4). The sizes of different regions were defined by preliminary experiments to optimally capture morphological changes during cell division and reflect the magnitude of change in morphology during the phases.

An ensemble model consisting of two deep learning models was used to detect mitotic subsections. The first model was a pretrained Inception v3 convolutional neural network. The head layers of the network were modified to output two values that are considered regression plane coordinates. The second model was a pretrained ResNet-50 network that is modified and trained for classification. Both models were pretrained on the ImageNet database. The ensemble model used both models for prediction. The prediction of the regression model was converted to a class and compared to the result of the classification model. If the two result classes were close to each other (their difference was less than a given threshold) then the ensemble model used the coordinates of the prediction model for cell selection, otherwise, it disregarded the predictions, and the corresponding cell was not selected for isolation.

To train the classification model, and to compare the predictions of the two models in the ensemble, regression plane coordinates were converted to discrete classes as follows. A rotation from the center of the regression plane was computed for each coordinate. The computation was performed according to the annotation strategy, i.e., 270 degrees in the Euclidean space is taken as 0 degrees rotation, and a clockwise (CW) direction is used. Coordinates between +/-15 degrees were assigned to the ‘interphase’ class. The remaining 330 degrees were equally split into 40 regions so that each region was 8.25 degrees wide. Interphase was indexed as 0 and each consecutive region in the CW direction got one higher index. The distance between classes could be computed using the indexing above, accounting for the circularity of the indices.

The dataset was the same for training the regression and classification models, however, for classification, the coordinates were converted to classes, as described above. The dataset was split into training, validation, and test sets in two steps. First, the validation set was separated by analyzing the trajectories of the cells in the regression plane. Images of the same trajectory (i.e. of the same cell) could only be split into different sets if their distance in the regression plane was at least 15 degrees. This approach ensured that very similar images would not be present in different sets, although this may result in empty regions of the regression plane in the validation set. This is especially an issue for prophase images. In the second step, the remaining dataset was split into training and test sets but without analyzing the trajectories. Instead, the density distribution of the 41 classes was analyzed and 3, 15, or 30 images were moved to the test set, depending on the number of elements in the class. This was required to have at least a few samples in each class in the test set.

The image intensities in the complete set were adjusted such that values between the lowest and highest 0.5% were mapped to values between 0 and 1. The training set was augmented before the training process using random histogram equalization using 64 bins, further histogram stretching, gaussian blur, and noise (Gaussian, speckle, or salt and pepper). Here, the density distribution was analyzed again and each section of 3 degrees was augmented to contain 2000 images in total. Further augmentation was applied during the training using random Gaussian noise, rotation between -90 and 90, scaling between 0.9 and 1.1, and translation of 3 pixels in both directions.

The training was performed in Matlab using two NVidia GeForce 2080 Ti 11 GB cards. A constant learning rate of 0.0001 was empirically set. Due to the large number of images in the augmented dataset, 5 epochs were sufficient for the training. The mini-batch size was set to 64. For evaluation during training, root mean square error (RMSE) was used for the regression model and accuracy for the classification model. If the class difference between the two prediction values was higher than 3 then the ensemble model discarded the sample and did not give a prediction. The ensemble model was evaluated on the test set and provided an RMSE of 679 on the 10K-by-10K regression plane while giving a prediction value for 97.5% of the samples. The regression model itself had an RMSE of 696. The classification model had an accuracy of 54.19% and a 3-miss accuracy of 95.74%. The performance could be further increased by keeping the samples that has a predicted radius value in the range of 3500 and 5000 pixels.

### UMAP projection of the datasets

To verify there is no domain shift between the distribution of the mitotic data set used for training and the predicted and selected image set, we extracted the final average pooling layer (2048-dimensional) from a ResNet-50 model trained for ensemble regression. The extracted features were first reduced to 16 dimensions using Principal Component Analysis (PCA) to mitigate noise. Uniform Manifold Approximation and Projection (UMAP) were applied. The UMAP transformation was fitted on the training subset of the mitotic dataset, and the reducer was applied to all dataset splits (training, validation, and test, and the atlas).

### Laser Microdissection

A Leica LMD6 system controlled by the Leica Laser Microdissection V 8.2.3.7603, equipped with a HC PL FLUOTAR L 63x/0.70 CORR XT objective was used for laser microdissection to prepare samples for mass spectrometry. Samples were dried thoroughly before the process. Registration of the fields of view of the Operetta screening microscope and LMD6 was performed using marker coordinates engraved in the PET membrane to allow easier detection of the selected cells. Fine alignment of the fields was performed with the SuperCUT unsupervised registration method^84^. 120 cells per class predicted by the ensemble model and visually confirmed by an expert, were dissected and collected in 16×24 cell culture plates (Greiner).

### Sample preparation for LC-MS analysis

Isolated cells were prepared for LC-MS analysis using an adjusted Deep Visual Proteomics workflow^16^. After laser microdissection 20 µl of 25 mM ammonium bicarbonate (ABC) buffer was added to the wells and samples were stored at -80 °C. For the sample preparation, the ABC was removed by lyophilization at 60 °C for 90 minutes. 4 µl of 60 mM triethylammonium bicarbonate (TEAB) was added to the wells. The plate was sealed using the Hamilton PCR ComfortLid and lysis was performed at 95 °C for 30 minutes. Afterwards, 1 µl of 60 mM TEAB in 60 % acetonitrile (ACN) was added and lysis continued at 75 °C for another 30 minutes. 1 µl of 4 ng/µl LysC was added, and the sample was digested at 37 °C for 3 hours. Then, 1.5 µl of 4 ng/µl trypsin was added and incubated at 37 °C overnight. The digestion was terminated by adding trifluoroacetic acid (TFA) to a final concentration of 1% v/v. Samples were lyophilized at 60 °C for 45 minutes. 4 µl of MS loading buffer (0.1% TFA in LC–MS-grade water) was added, and the plate was vortexed for 30 seconds and centrifuged at 2,000g for 5 minutes.

### Proteomic Measurements

LC–MS analysis was performed with an EASY-nLC-1200 system (Thermo Fisher Scientific) connected to the timsTOF Ultra (Bruker Daltonik) with a nano-electrospray ion source (CaptiveSpray, Bruker Daltonik). Peptides were loaded on a 50-cm in-house-packed HPLC column (75-µm inner diameter packed with 1.9-µm ReproSil-Pur C18-AQ silica beads, Dr. Maisch).

Peptides were separated using a linear gradient from 5–30% buffer B (0.1% formic acid and 80% ACN in LC–MS-grade water) in 55 minutes, followed by an increase to 60% for 5 minutes and a 10-minute wash in 95% buffer B at 300 nl min^−1^. Buffer A consisted of 0.1% formic acid in LC–MS-grade water. The total gradient length was 70 minutes, and the column temperature was 60 °C.

Mass spectrometric analysis was performed as previously described ^22^ using data-independent acquisition (diaPASEF) and high-sensitivity mode. The elution of precursors from each ion mobility scan was synchronized with the quadrupole isolation window. 16 windows were scanned in each PASEF cycle. The collision energy was ramped linearly as a function of the ion mobility from 59 eV at 1/K0 = 1.6 Vs cm−2 to 20 eV at 1/K0 = 0.6 Vs cm−2.

### Data analysis of proteomic raw files

Raw files were analyzed with the neural network-based search engine DIA-NN (version 1.8.1)^85^ against the UniProt database (March 2023 release, UP000005640_9606 and UP000005640_9606_additional). The search was performed library-free. Fragment and precursor m/z were set to a minimum of 100 m/z and a maximum of 1700 m/z. Peptides were allowed to be between 7 and 30 amino acids long. The precursor charge range was set from 2 to 4. The maximum number of missed cleavages and maximum number of variable modifications was set to 2. N-terminal methionine excision was enabled, and methionine oxidation and N-terminal acetylation were set as variable modifications. Mass accuracy was fixed to 1.5e-05 (MS2) and 1.5e-05 (MS1). Match between runs was enabled. 2 empty raw files were excluded from the search.

### Processing and visualization of proteomic data

Out of 123 analyzed samples, three samples with less than 3000 protein groups were excluded to avoid statistical issues after imputation. 1385 of 5735 protein groups showed fewer than 67% quantification across the samples of at least one replicate and were removed using Perseus (v2.0.5.0)^86^. Batch correction on replicates was applied using ComBat^87^. AlphaPeptStats^88^ was used to remove contaminants. Differential abundance analysis, Kruskal-Wallis test between mitotic classes, and Z-scoring was performed in Perseus^82^. 147 significantly changed proteins were identified applying the Kruskal-Wallis test on the 41 subsections with a Benjamini-Hochberg q-value of less than 0.1. To increase statistical power, three consecutive mitotic subsections were grouped, e.g., 1, 2-4, 5-7, …, 39-41. Interphase remained as a single subsection. Subsection 20 was grouped with subsections 17, 18, and 19. A Kruskal-Wallis test was performed on these 14 grouped subsections resulting in 1060 proteins that were significantly changed with a Benjamini-Hochberg q-value of less than 0.05. Data was visualized using the numpy (v1.24.3)^89^, pandas (v2.0.3)^90^, matplotlib(v3.7.1)^91^, and seaborn (v0.12.2)^92^ packages in Python (v3.10.11), as well as ggplot2 (v3.4.2)^93^ and ComplexHeatmap (v2.16.0)^94^ packages in R (v4.3.0) using RStudio (build 524).

### Hit selection

List of proteins whose expression significantly changes during the process of mitosis were detected with the Kruskal-Wallis statistical test at false discovery rate (FDR) 0.1 and 0.05 (with Benjamini-Hochberg correction). Prior to running the test, the proteomic data was Z-scored. This analysis was performed with Perseus software.

### Network analysis of proteomic data

To identify interactions between the most relevant proteins, the Hugo IDs of proteins changing significantly during cell cycle were used as input for the STRING^47^ v12.0 online database. We selected interactions having at least medium confidence score, encompassing both experimentally verified and predicted interactions. The resulting network was exported and further analyzed using the Cytoscape^48^ software v3.9.1.

Clusters in the protein-protein interaction network were identified using the MCL clustering algorithm in the AutoAnnotate^95^ v1.4.0. Cytoscape plugin. Clusters were named by using the gene function showing the strongest enrichment in the GOrilla^96^ Gene Ontology enRIchment anaLysis and visuaLizAtion tool.

To assess the strictness of expression regulation during mitosis, we scaled the median expression values of each protein in the 41 subsections of the cell cycle and subsequently fitted a 3rd order polynomial regression model. Then, the sum of the absolute values of residuals was calculated for each protein.

### Analysis of cancer driver genes

The multi-omics dataset of driver mutations was downloaded from DriverDBv4 (Ver. 1.00.003). Data on mutations in 32 tumor types in The Cancer Genome Atlas (TCGA) database were downloaded from cBioPortal (https://www.cbioportal.org/, downloaded on 07/11/2023). We used the Cancer Bayesian Selection Estimation (CBaSE) v1.1 tool^54^ to acquire gene-specific probabilistic estimates of the strength of positive selection in cancer samples belonging to each tumor type. For the final analysis only those tumor types were selected where at least 20,000 missense mutations were present (see Fig. 3e). The COAD and READ were jointly deposited in the cBioPortal database. Therefore we refer to both of them as COAD/READ in the text. We examined genes encoding proteins that were identified in the proteomics analysis. We ran cBaSE with default parameters and considered positively selected genes with a q value lower than 0.05.

### Statistics and Figure Preparation

Sample sizes were similar to those generally applied in the field. Data was tested for normality using a Shapiro-Wilk test in order to determine the appropriate statistical method to use. Data met the necessary criteria for all analyses used.

Figure preparation was performed with the Adobe Photoshop (23.2.1) and Illustrator (27.4.1) programs. Images presented on the same figure were modified identically.

### Code and data availability

Mitotic image and proteomics dataset is available at https://mito-omix.org/ Codes used for the data analysis is available at: https://github.com/biomag-lab/DVP2

The mass spectrometry proteomics data have been deposited to the ProteomeXchange Consortium via the PRIDE partner repository^97^ with the dataset identifier PXD047018.

## Funding

We acknowledge support from the Lendület BIOMAG grant (no. 2018–342), TKP2021-EGA09, Horizon-BIALYMPH, Horizon-SYMMETRY, Horizon-SWEEPICS, H2020-Fair-CHARM, CZI Deep Visual Proteomics, HAS-NAP3, the ELKH-Excellence grant from OTKA-SNN no. 139455/ARRS, the FIMM High Content Imaging and Analysis Unit (FIMM-HCA; HiLIFE-HELMI), and Finnish Cancer Society. MM, AM, and FP are funded by the Novo Nordisk Foundation (grant agreements NNF14CC0001 and NNF15CC0001). MM is funded by the Max Planck Society for the Advancement of Science. FP is additionally funded by the Novo Nordisk Foundation (grant agreement NNF0069780). We acknowledge Ulrike Kutay (ETH, Zurich, Switzerland), Zoltan Lipinszki (BRC, Szeged, Hungary) and Farkas Sukosd (University of Szeged, Hungary) for the constructive discussions about the results.The results shown here are partly based upon data generated by the TCGA Research Network: https://www.cancer.gov/tcga.

## Author contributions

Conceptualization: EM, VM, FP, CP, AM, EL, MM, PH

Methodology: EM, VM, FP, KK, AB, DK, NH, IG, NM, RH, FK, AK, MM, AM, MM, PH

Investigation: EM, VM, FP, ZZI, RH, DC, FK, UA

Visualization: EM, VM, FP, DK, ZZI, IG, NM, RH, MM, PH

Funding acquisition: EL, MM, PH

Project administration: VM

Supervision: CP, EL, MM, AM, MM, PH

Writing – original draft: EM, VM, FP, NM, AM, EL, MM, PH

Writing – review & editing:

### Competing interests

P.H. is the founder and a shareholder of Single-Cell Technologies Ltd., a biodata analysis company that owns and develops the BIAS software. The remaining authors declare no competing interests.

## Supplementary Materials

### Extended data figures

**Fig S1.**
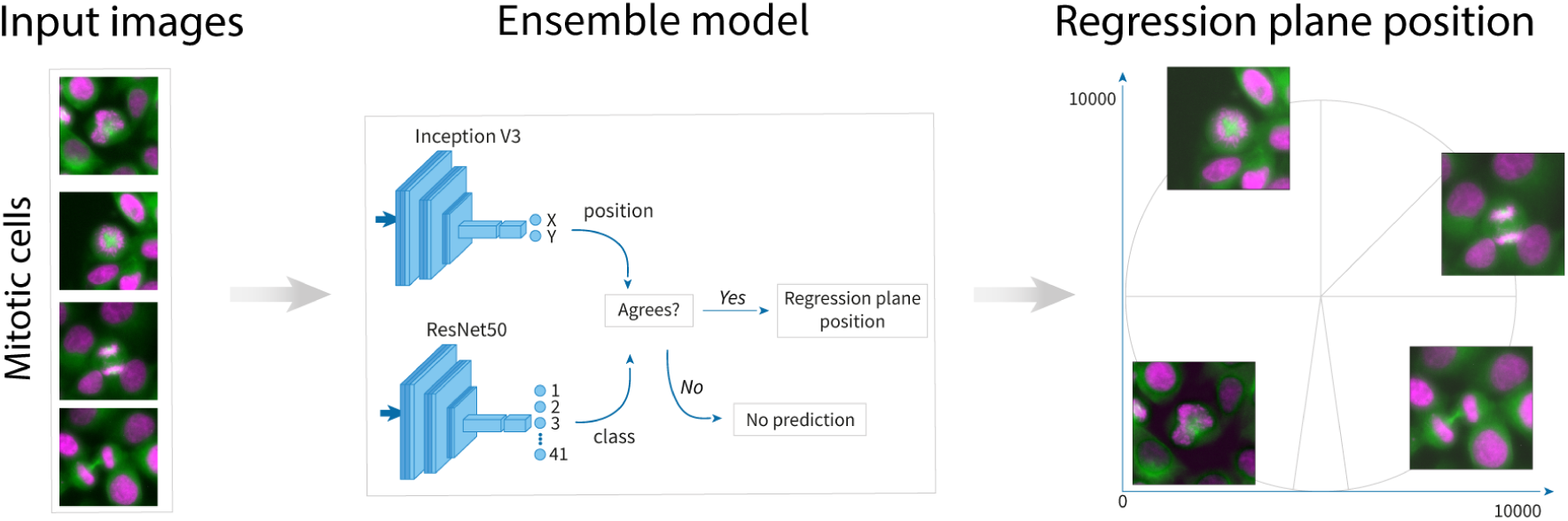
Architecture of the ensemble model for mitotic subsection prediction. Image crops of cells were fed into an ensemble model consisting of two deep learning models to predict their mitotic subsections. The first model was a pretrained Inception v3 convolutional neural network with modified head layers to output two values that are considered regression plane coordinates. The second model was a pretrained ResNet-50 network trained for classification. The prediction of the regression model was converted to a class and compared to the result of the classification model. If the two result classes were close to each other then the ensemble model used the coordinates of the prediction model for cell selection, otherwise, it disregarded the predictions, and the corresponding cell was not selected for isolation.

**Fig S2.**
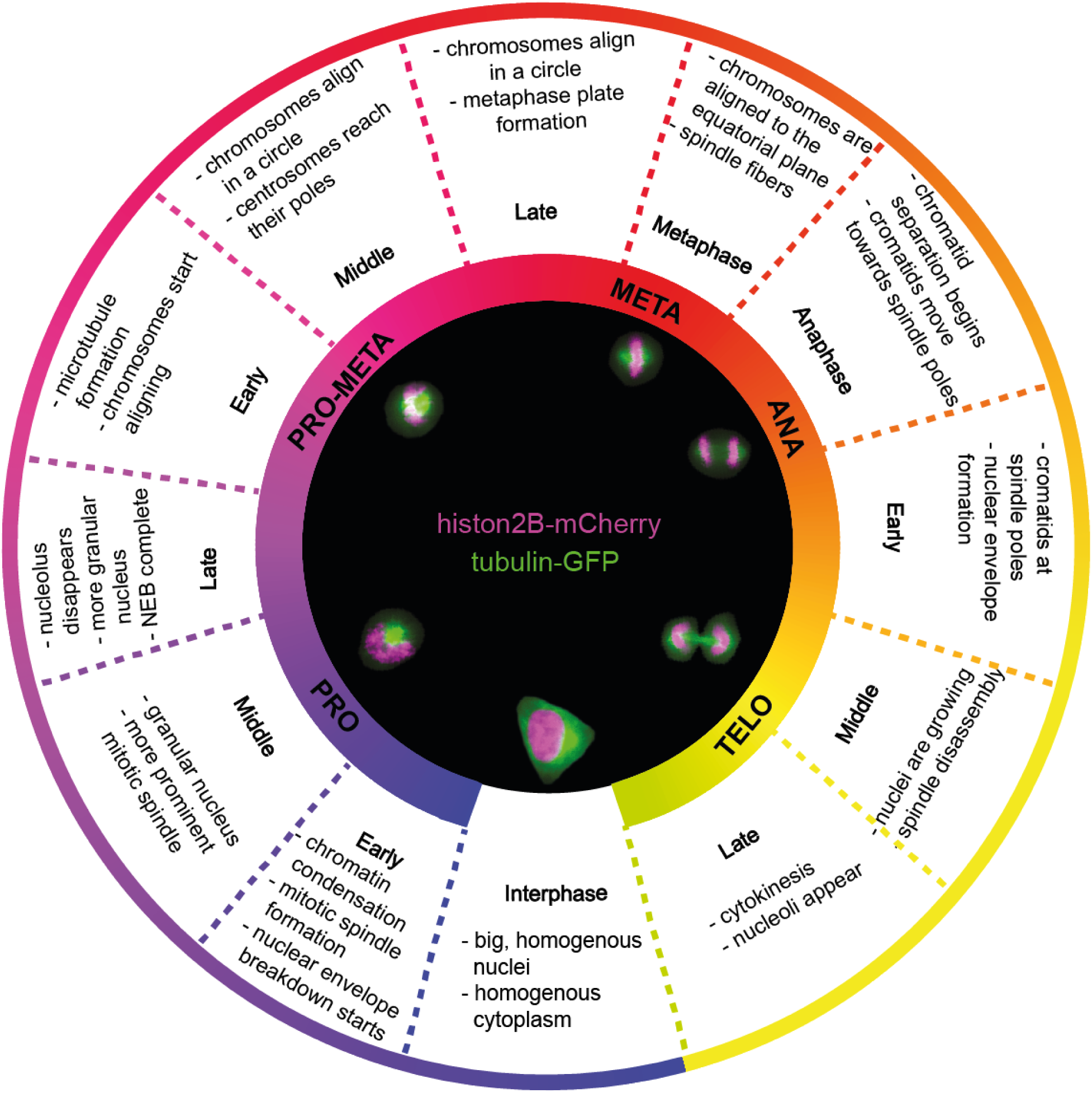
General guidelines for regression plane annotation. The inner circle shows example cells for each mitotic phase and interphase, the outer circle shows guidelines that were used for determining the approximate coordinates of the cells in the training set during the model building phase.

**Fig S3.**
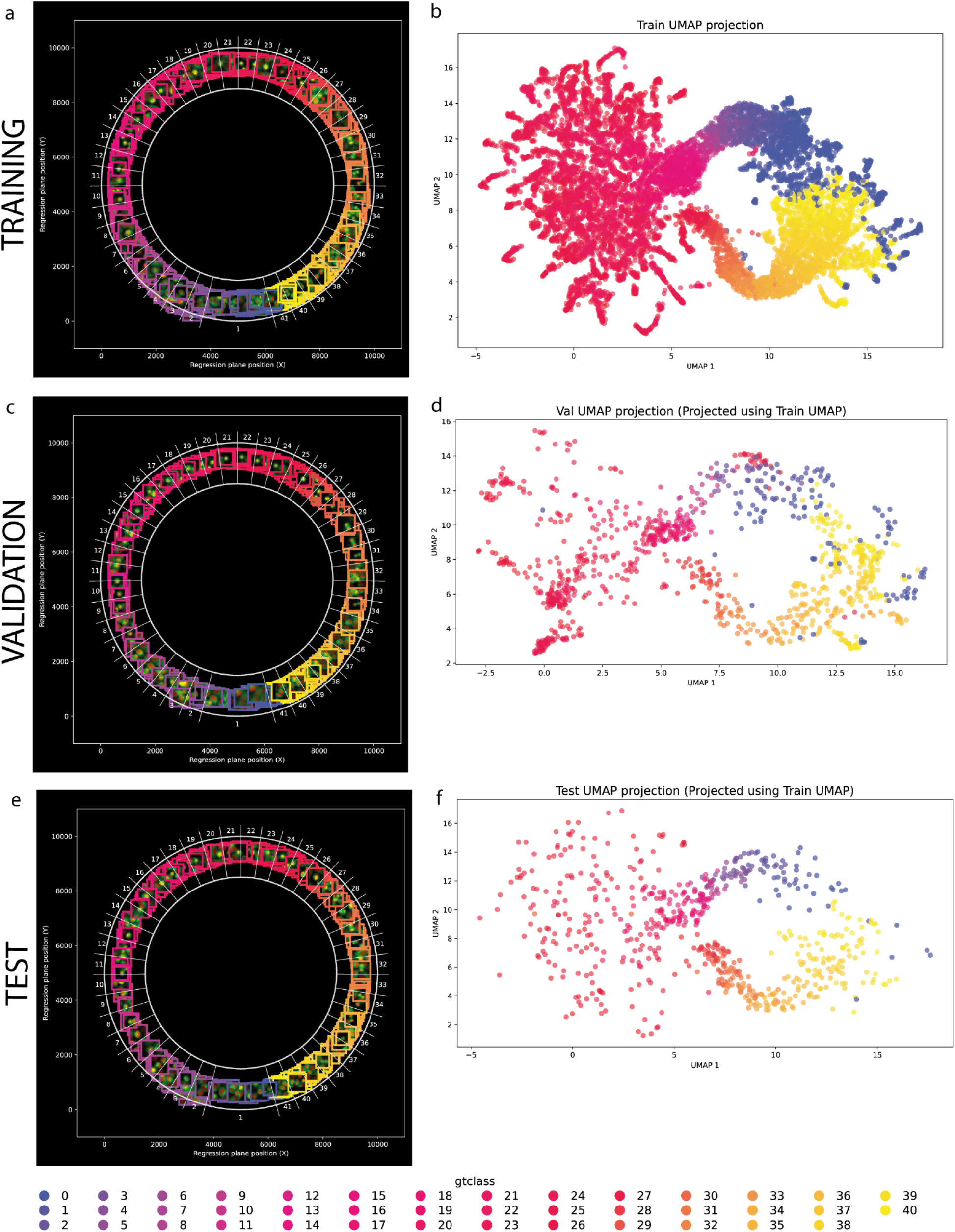
Visualization of the mitotic regression plane dataset. a-c-e shows the cells’ annotated position of the regression plane for the different training set. b-d-f Corresponding UMAP projection for each sets. UMAP’s reducer were computed using the training data subset, and each of the sets projected with the training reducer.

**Figure S4:**
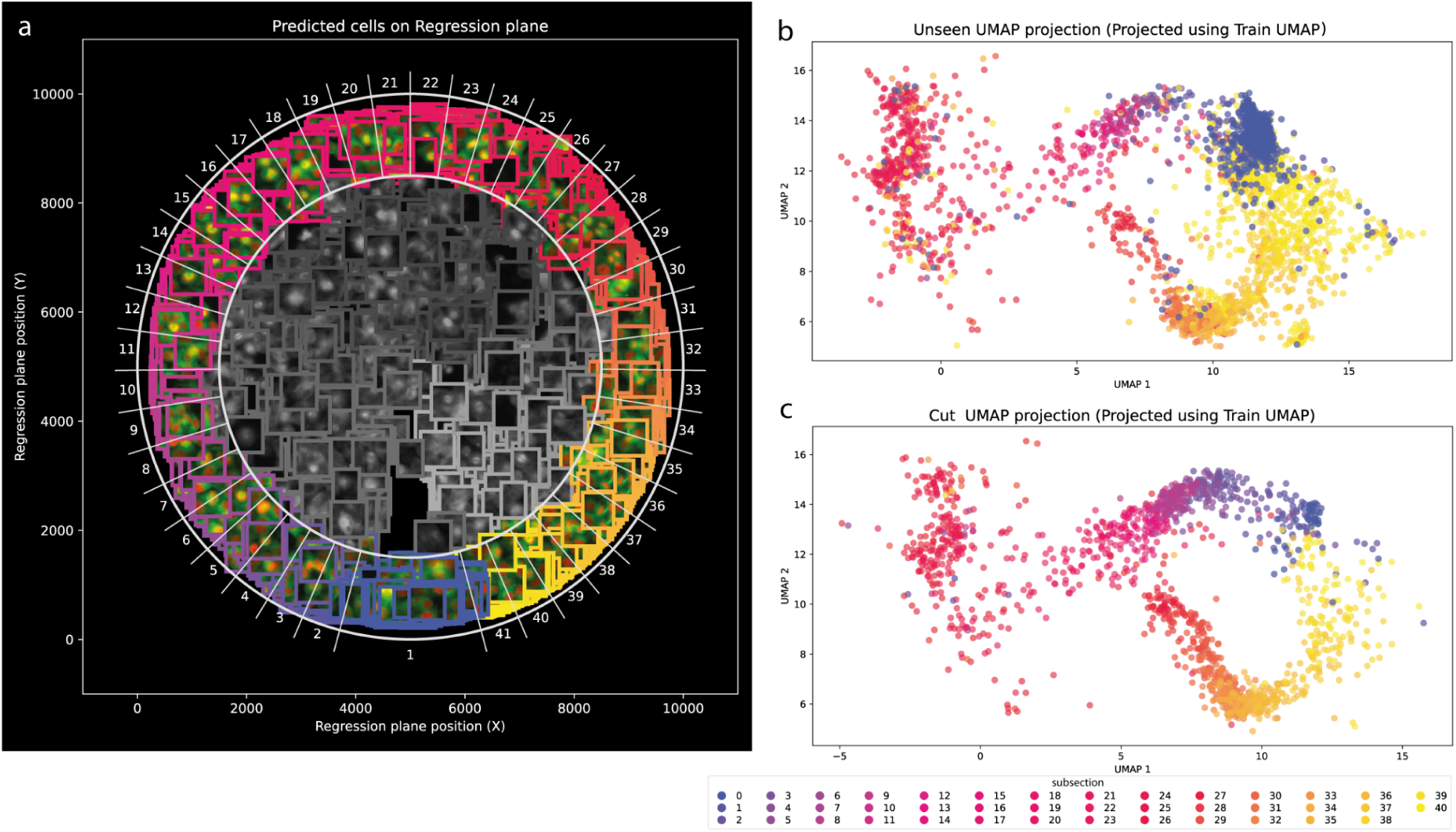
Visualization of the Predicted Cell Positions and UMAP Projections. a Predicted cell positions from a sample slide mapped onto the regression plane. Individual cells are visualized according to their position on the regression plane. Cells with a radius below 3500 were discarded by the model and are shown as gray images. b UMAP projection of the prediction set, including 3% interphase cells, generated using the UMAP reducer from the training data. c UMAP projection of the subset of predicted cells that were manually verified and selected for single cell cutting by an expert.

**Fig S5.**
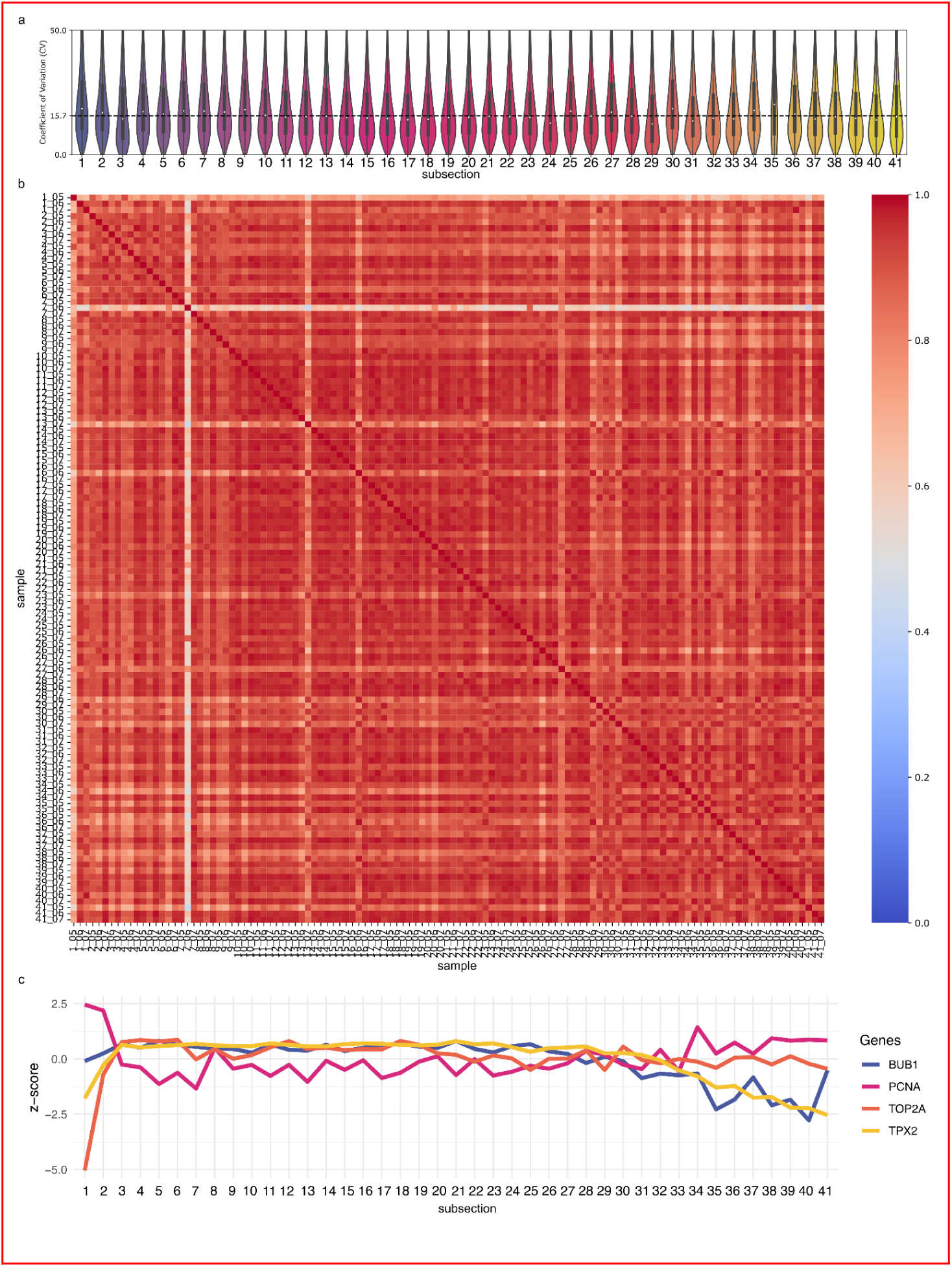
Data quality of proteomic measurements. a) Coefficient of variation of log2 protein intensities for each mitotic subsection. The dashed line shows the average median of the coefficient of variation across the subsections. The y-axis was limited at 50% to enhance visibility of median values. b) Pearson correlation matrix of raw protein intensities across all samples. c) Mean z-score trajectory of mitotic markers and DNA replication to validate the correct isolation of mitotic subsections. BUB1, PCNA, TOP2A, and TPX2 were selected as markers.

**Fig S6.**
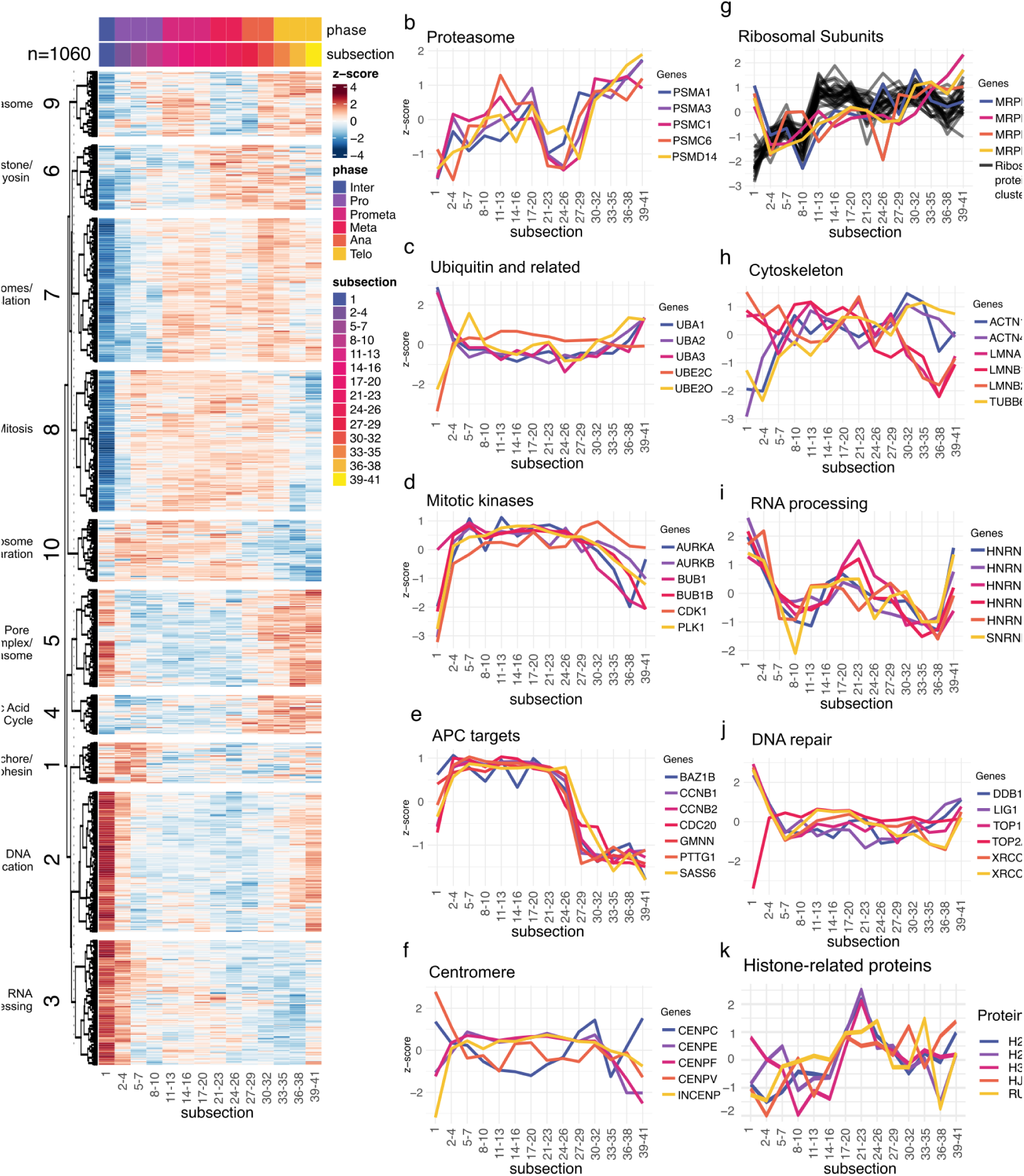
Proteome dynamics of 1060 significantly changed proteins across 14 subsections. a) Heatmap of significantly changed proteins (Kruskal-Wallis test with Benjamini-Hochberg q-value of 0.05) across 14 subsections. 41 subsections were regrouped resulting in 14 subsections with about 9 replicates per subsection. Protein abundance dynamics across 14 mitotic subsections of the proteasome (b), ubiquitin and related proteins (c), mitotic kinases (d), APC targets (e), centromere subunits (f), ribosomal subunits (g), cytoskeleton (h), RNA processing (i), DNA repair (j), histone-related proteins (k).

**Fig S7.**
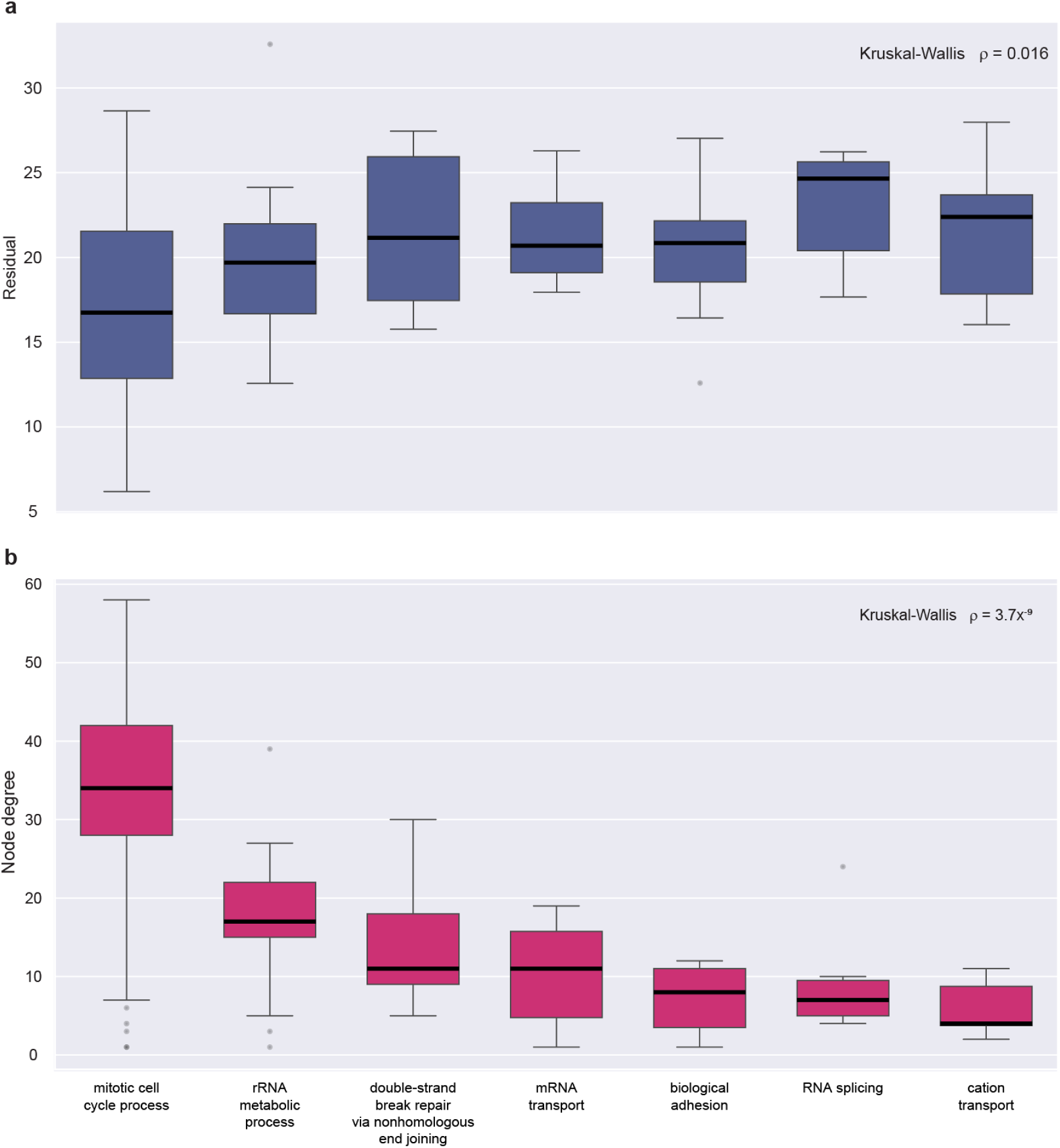
Node degree and sum of residuals in different functional clusters. The groups are ordered based on median node degree. Two-sided P value of Kruskal-Wallis tests are indicated.

**Fig S8.**
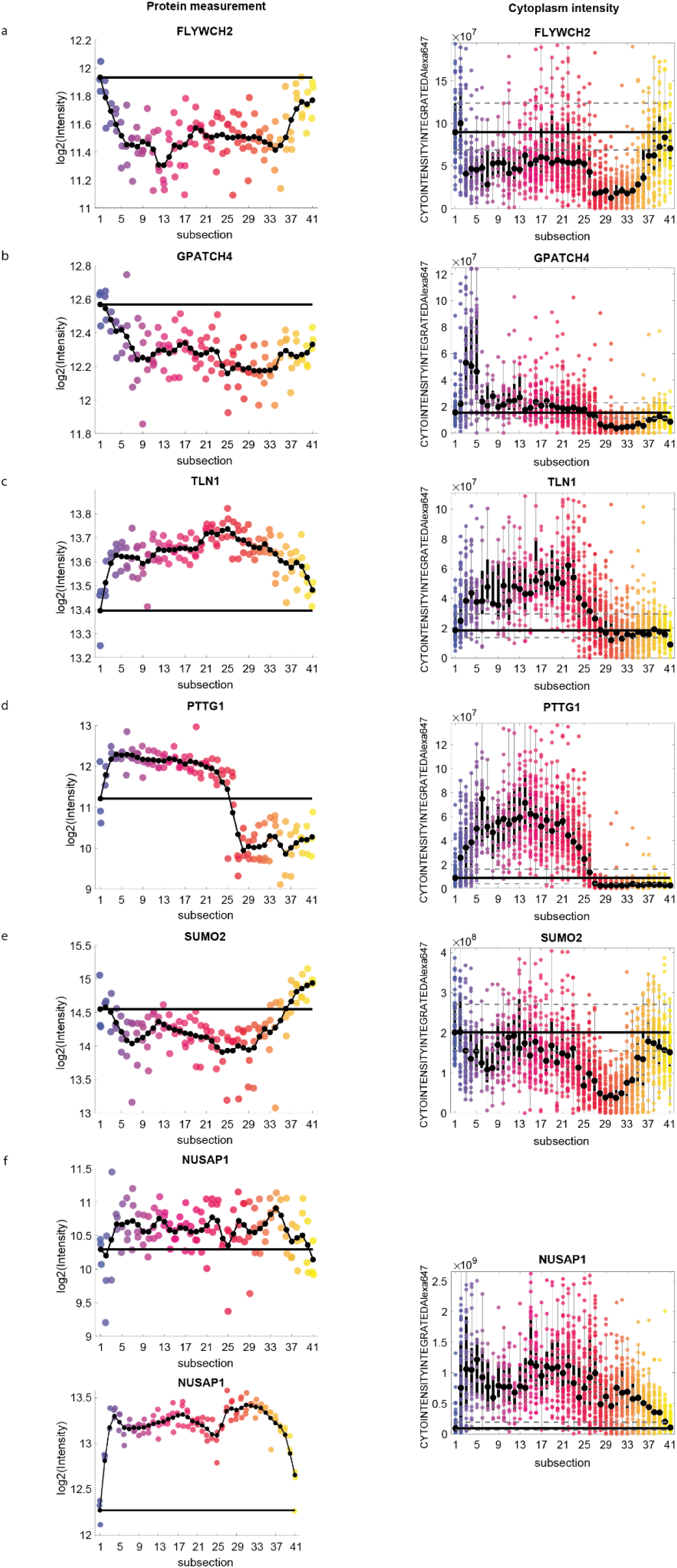
Proteomic measurement (left) and protein abundance estimated by immunofluorescence within the cell (right) for the respective proteins (Part 1). a) FLYWCH2 *r_s_* =− 0. 174 *P* = 0. 02 b) GPATCH4 *r_s_* =− 0. 33 *P* = 0. 004 c) TLN1 *r_s_* = 0. 782 *P* < 0. 001 d) PTTG1 *r_s_* = 0. 754 *P* < 0. 001 e) SUMO2 *r_s_* =− 0. 522 *P* < 0. 001 f) NUSAP1 *r_s_* = 0. 18 *P* < 0. 001 NUSAP1 *r_s_* = 0. 20*P* = 0. 006

**Fig S9.**
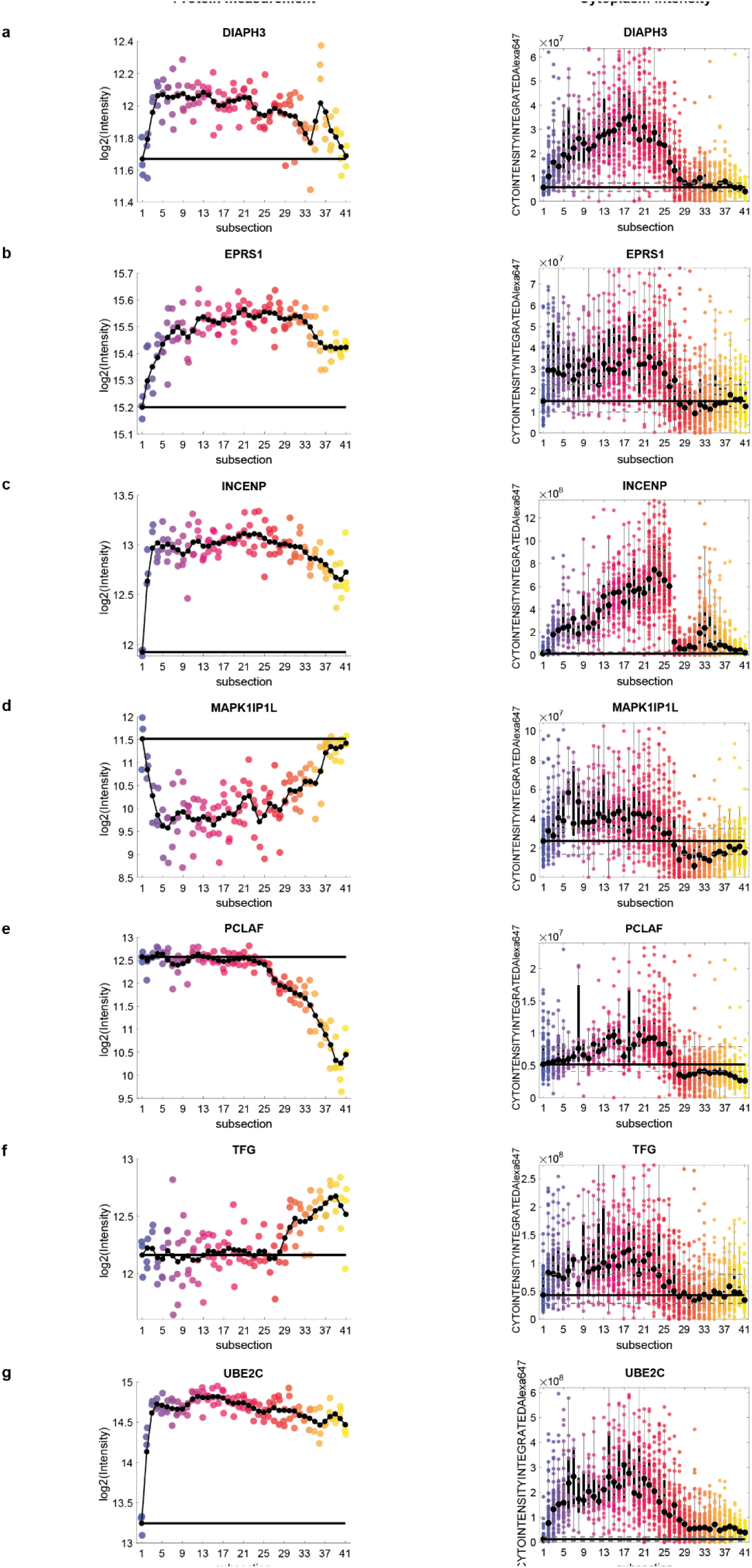
Proteomic measurement (left) and protein abundance estimated by immunofluorescence within the cell (right) for the respective proteins (Part 2). a) DIAPH3 *r_s_* = 0. 255 *P* = 0. 0013 *r_s_* = 0. 599 *P* < 0. 001 *r_s_* =− 0. 005 *P* = 0. 147 *r_s_* = 0. 447 *P* < 0. 001 b) EPRS1 d) MAPK1IP1L f) TFG *r_s_* = 0. 460 *P* < 0. 001 *r_s_* =− 0. 476 *P* < 0. 001 *r_s_* =− 0. 132 *P* = 0. 575 c) INCENP e) PCLAF g) UBE2C

**Extended Data Table 1.**
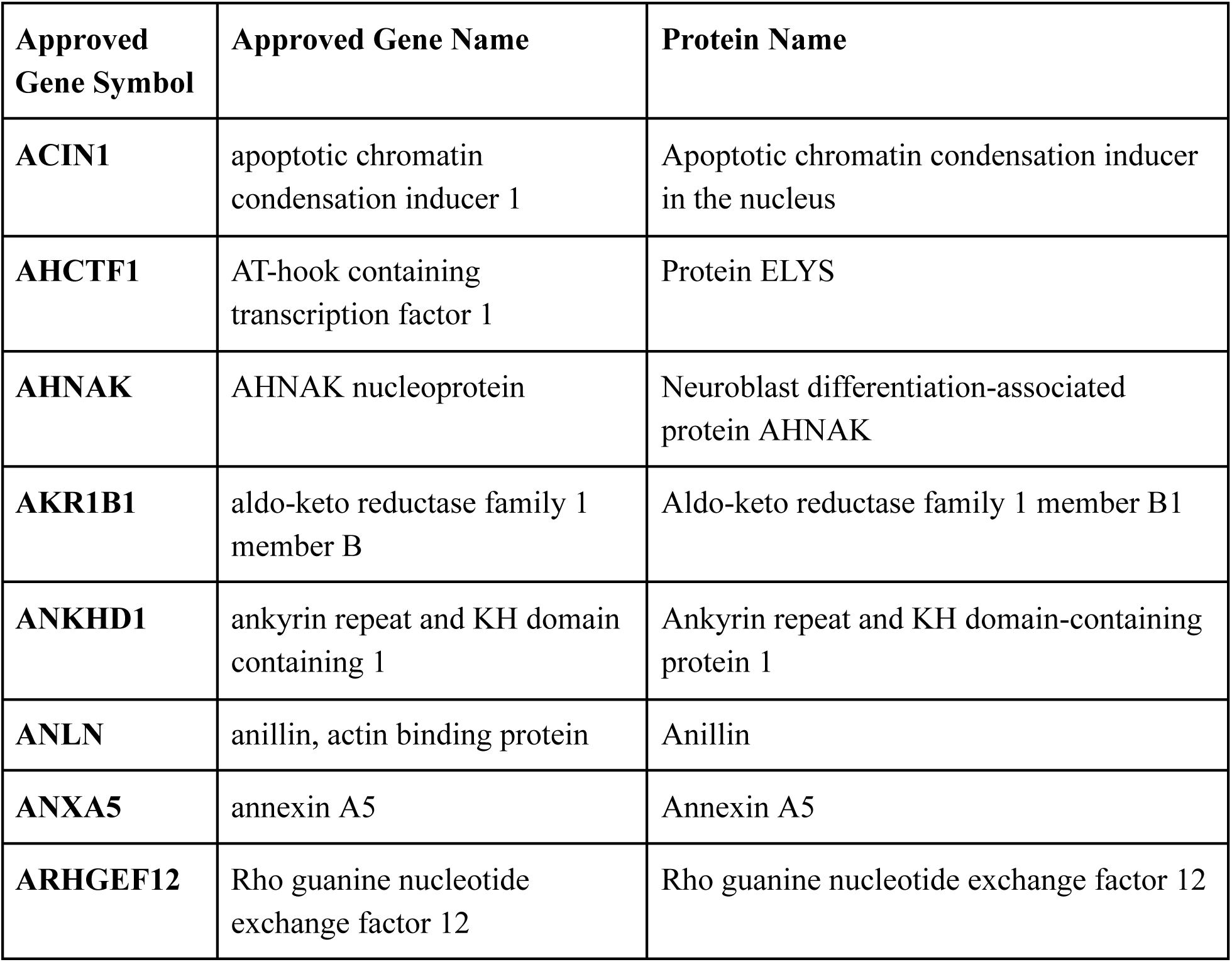
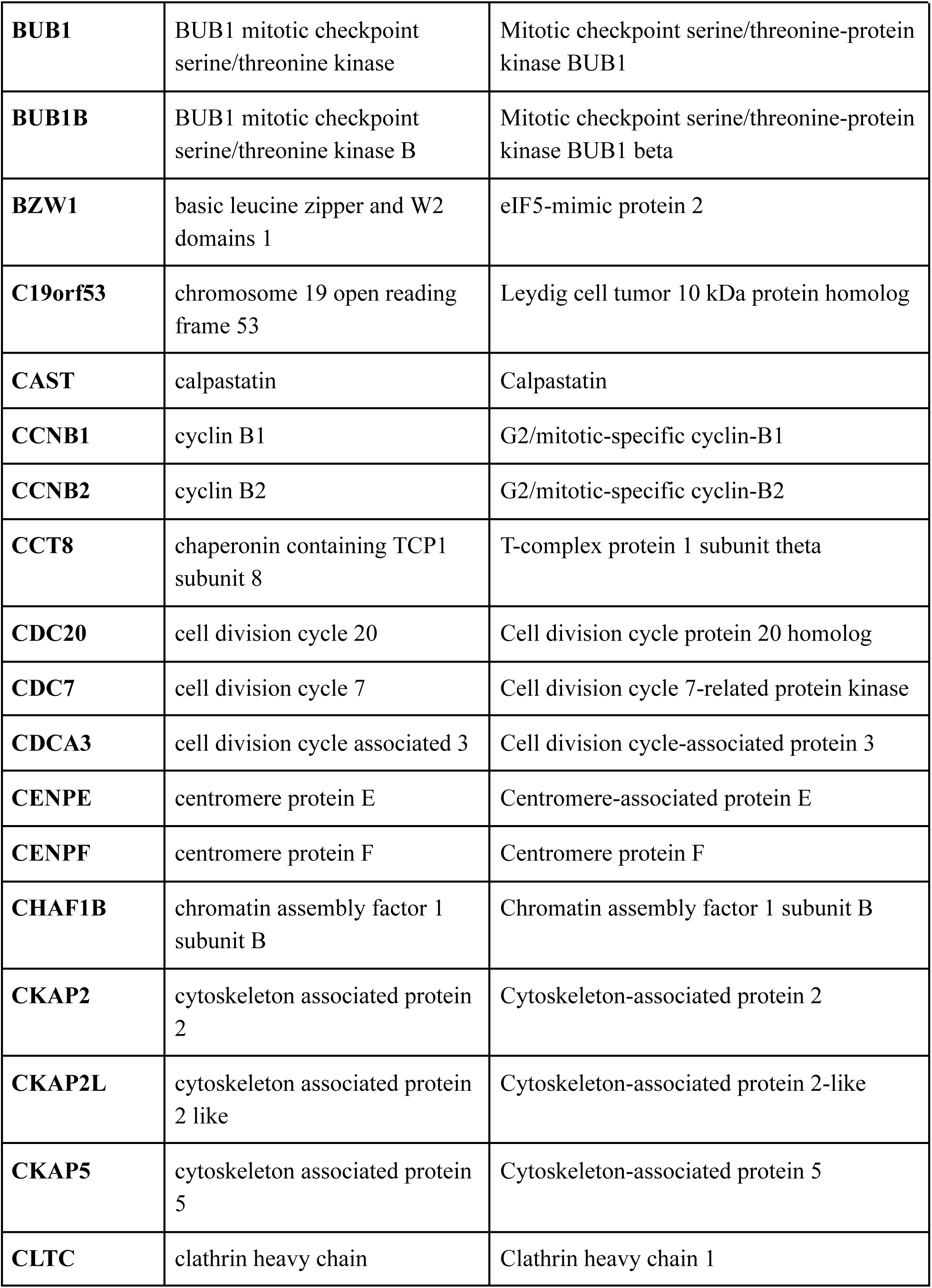
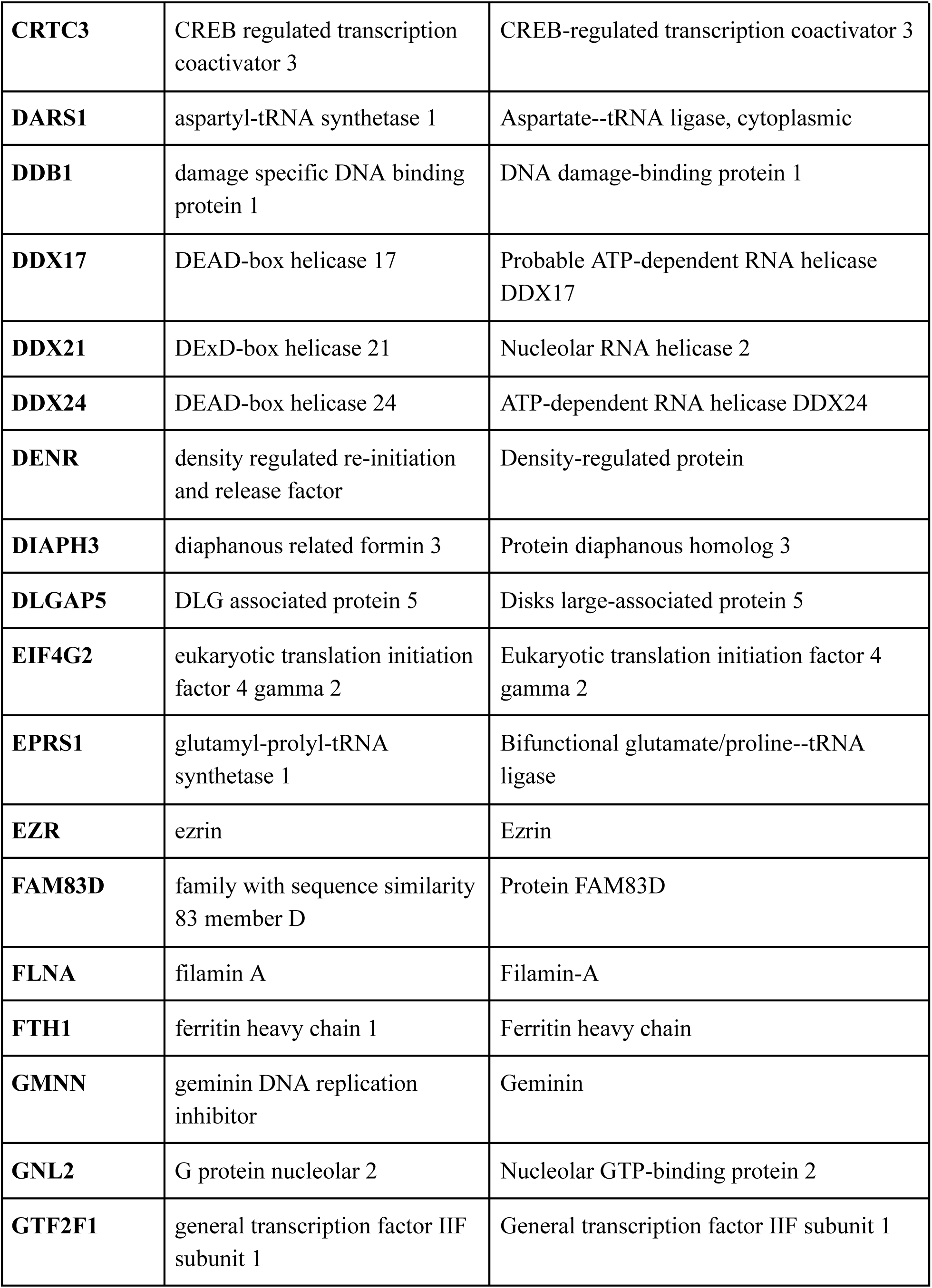
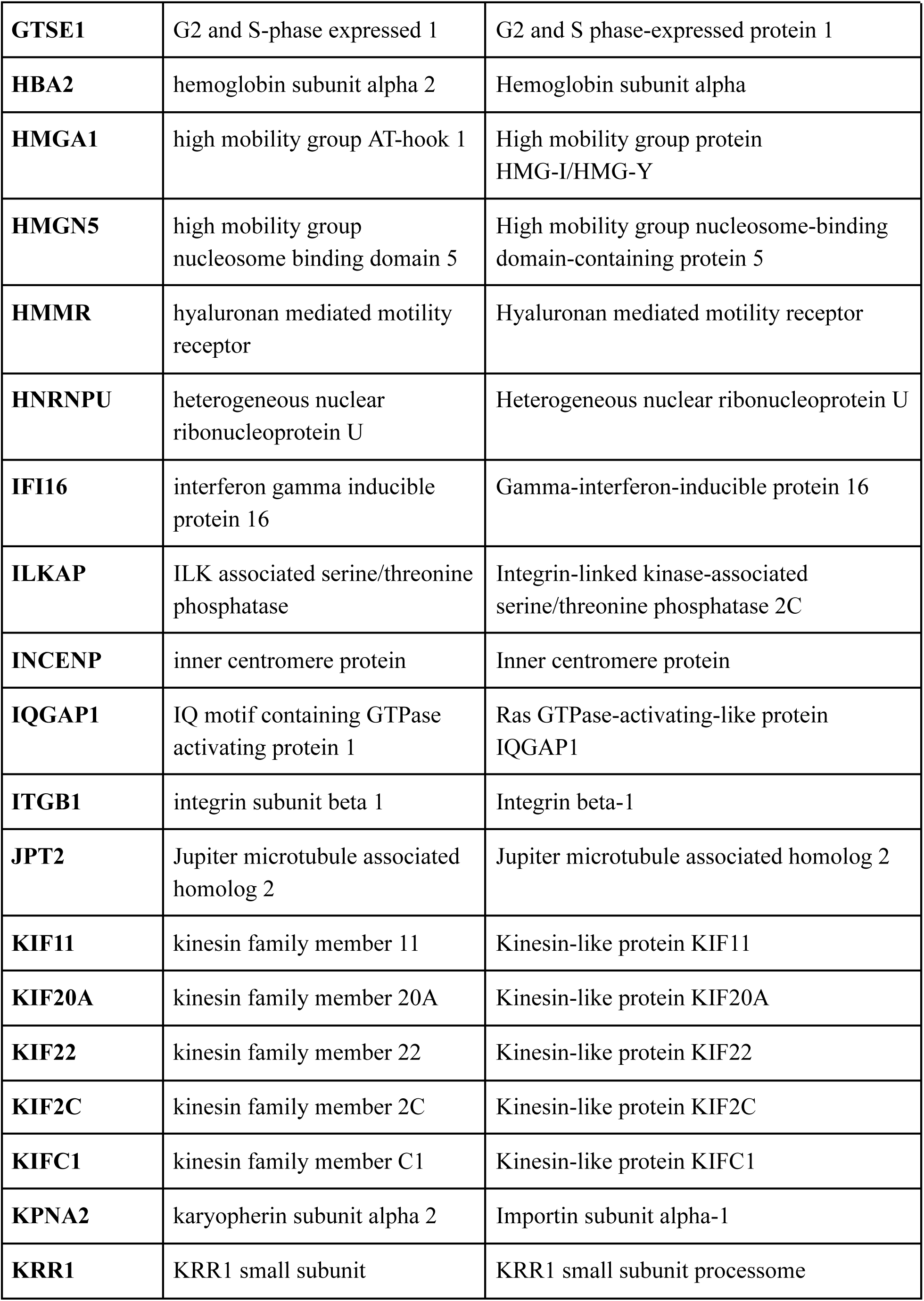
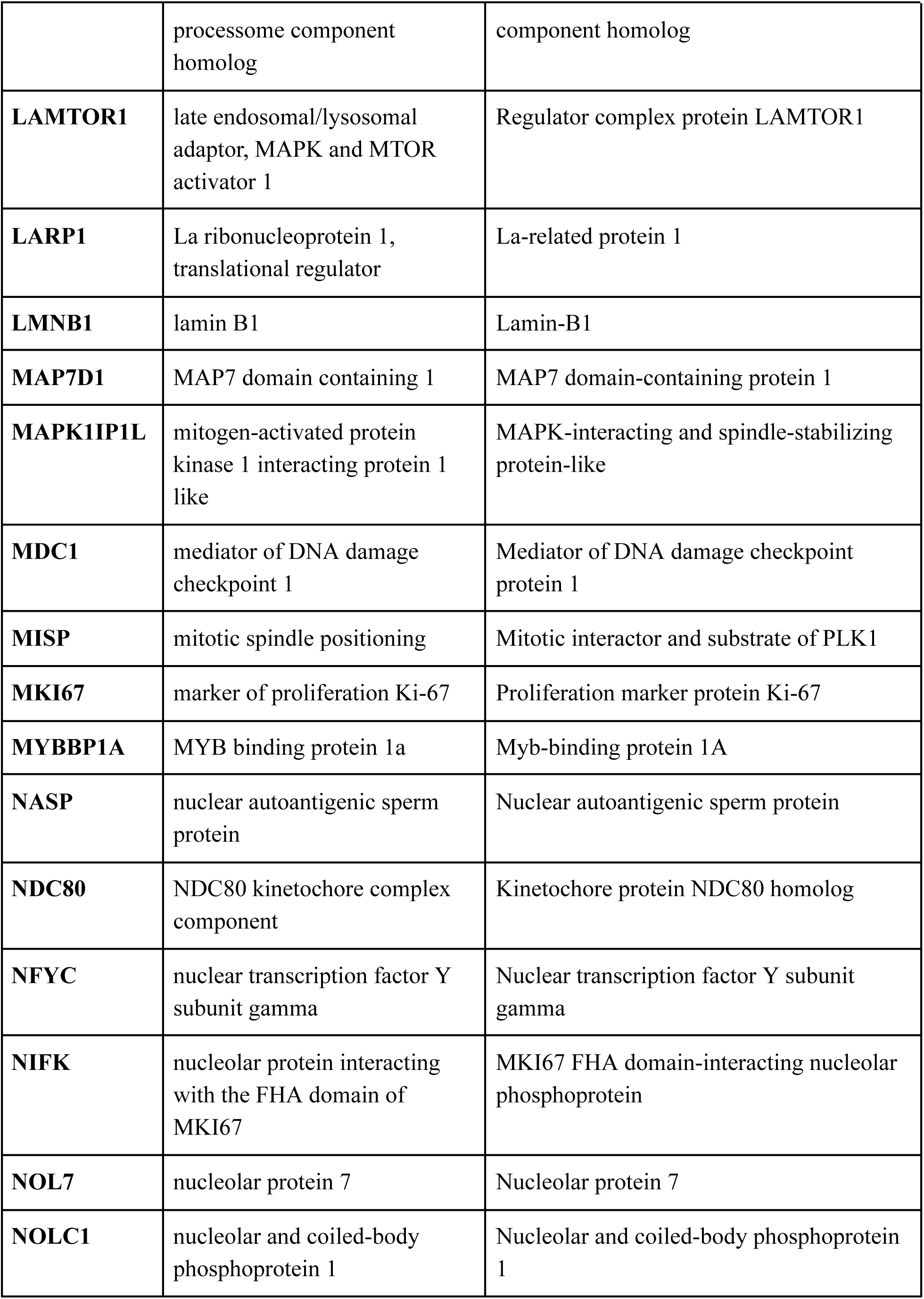
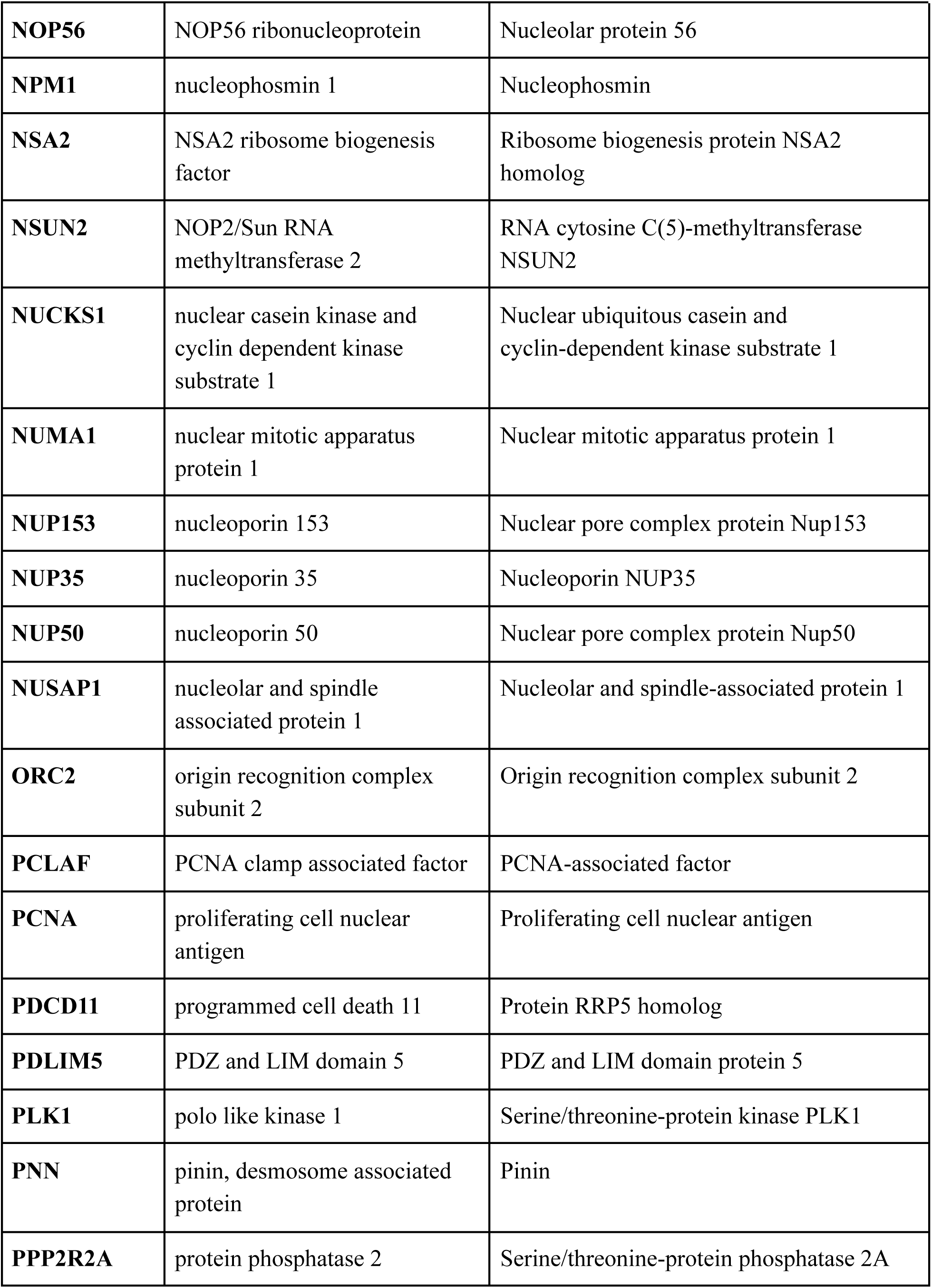
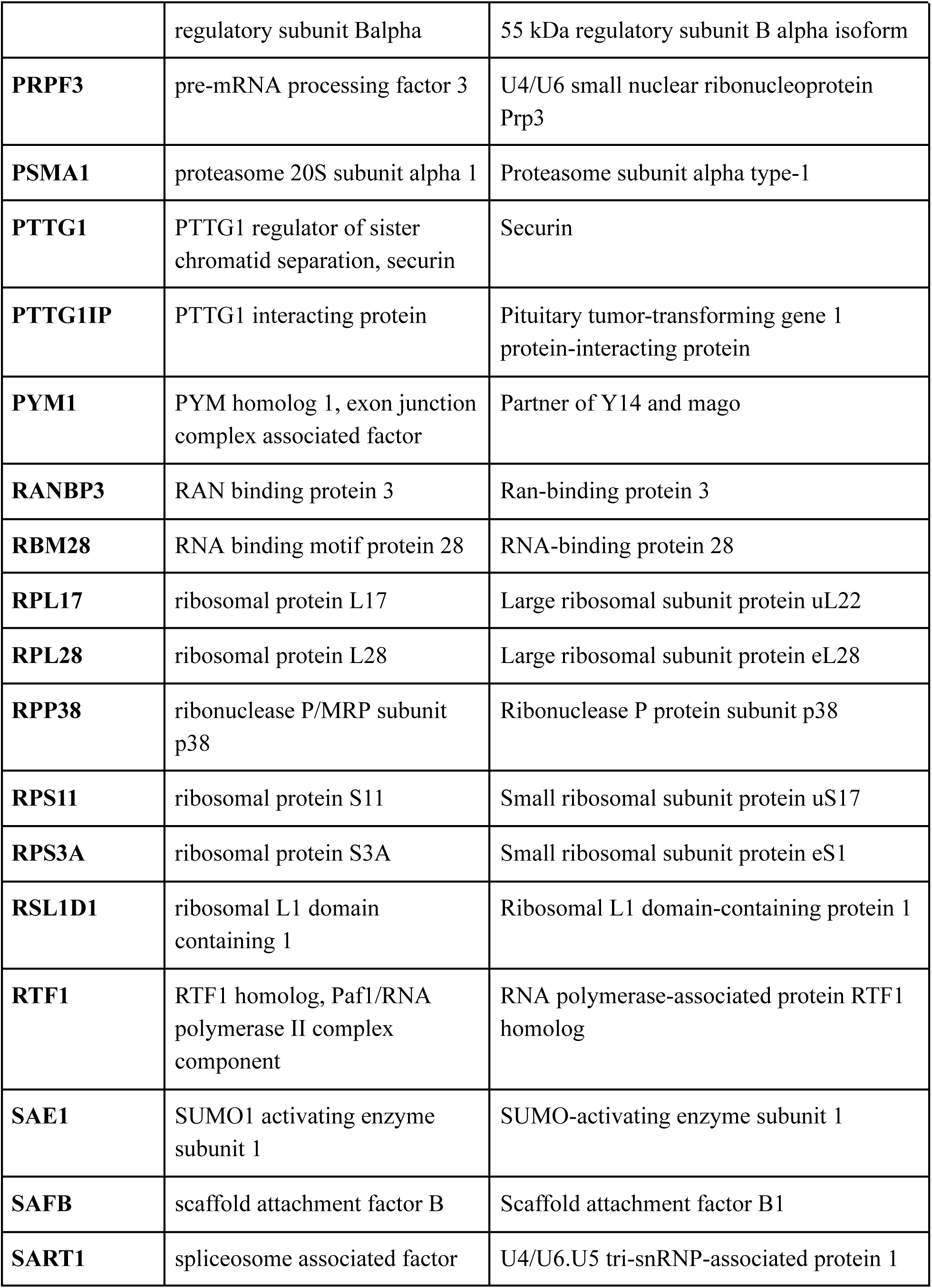
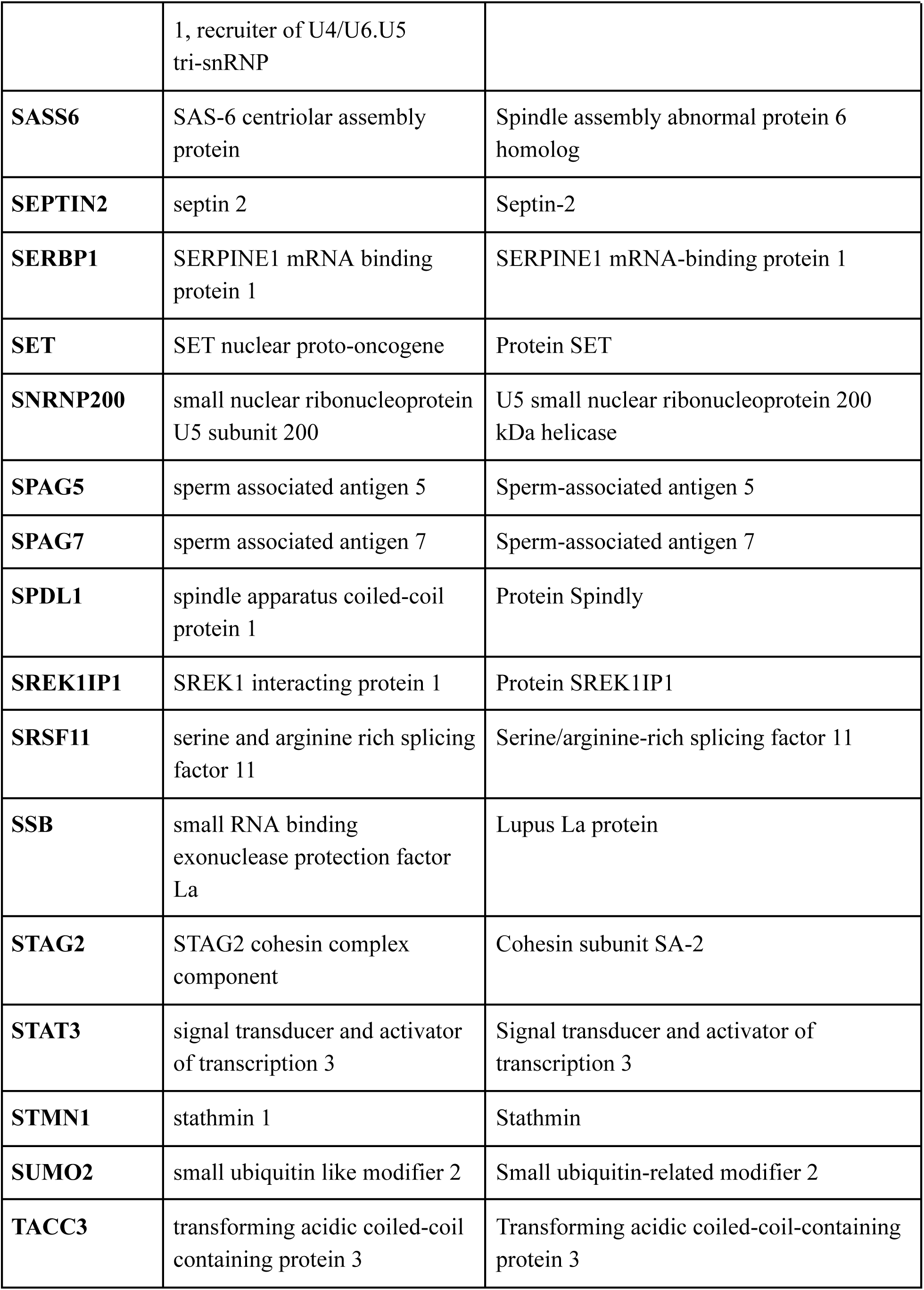
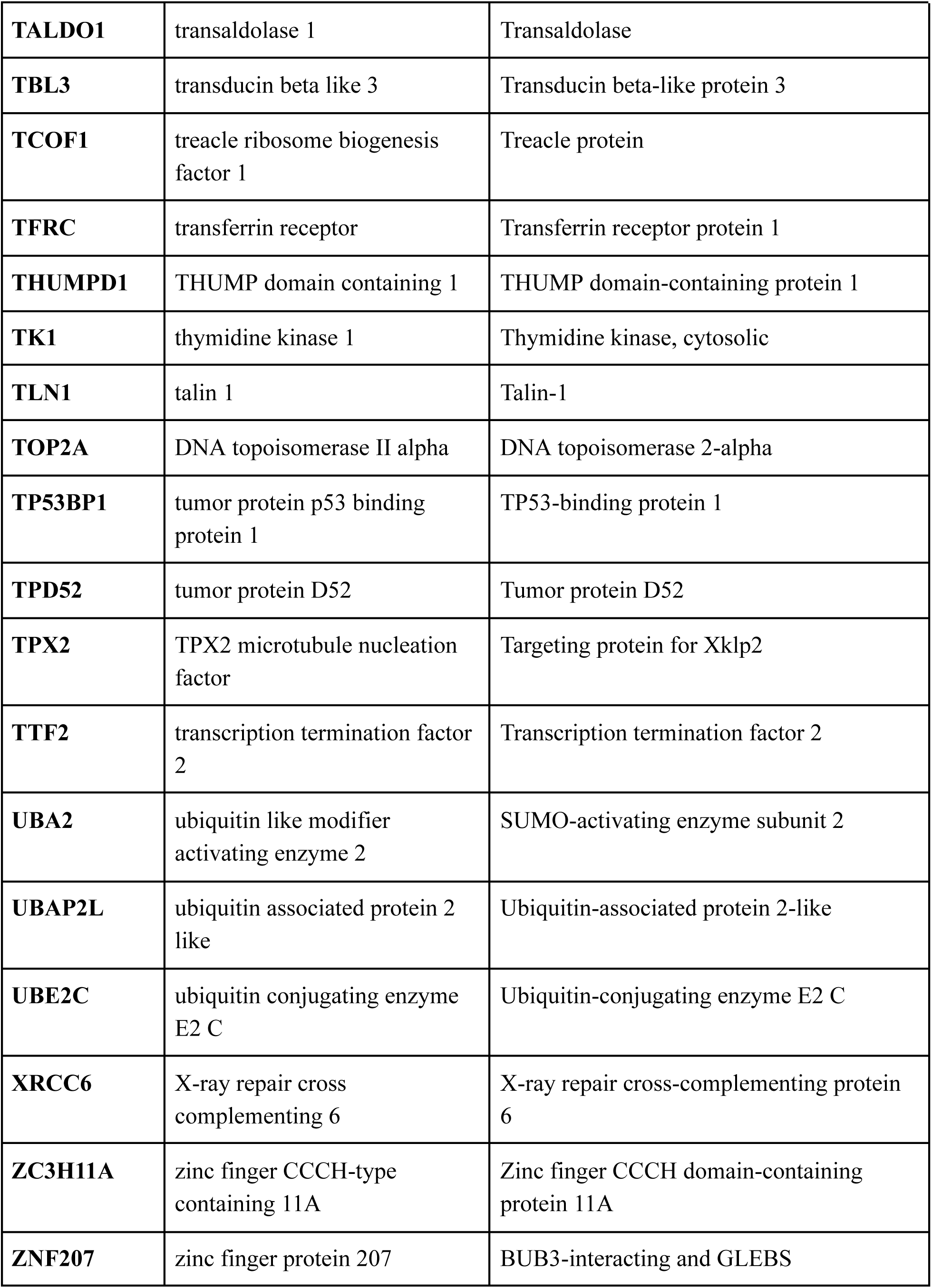
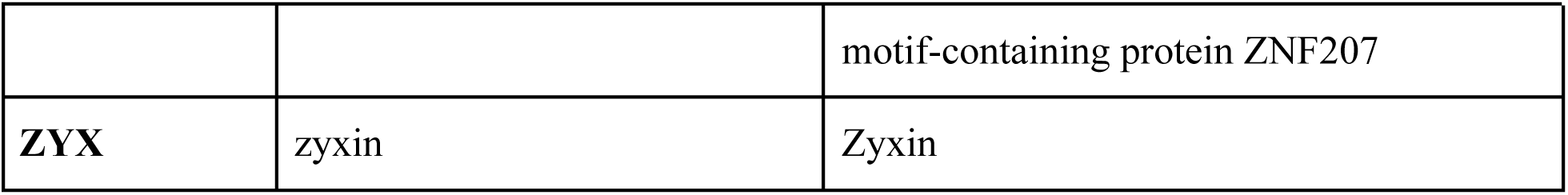
Gene names for the proteins with significantly changing abundance during mitosis.

**Extended Data Table 2.** The association between residuals and node degree remains consistent, irrespective of protein expression levels and biological replicates. We evaluated the influence of potential confounders, including median protein expression and biological replicates, using a multivariate linear regression model. The significance of node degree remained significant. The effect of replicates B and C were compared with replicate A. The P-value reflects the likelihood that the observed relationship between the predictor and response variables occurred by chance. R² represents the proportion of total variance in residuals explained by the model, adjusted for the number of variables. A non-significant P-value from the Breusch-Pagan (BP) test suggests the absence of heteroscedasticity.

| Variable | Estimate | p-value |
| --- | --- | --- |
| Node degree | -0.1471 | $<2.2 \times 10^{-16}$ |
| Median expression | 0.3728 | 0.004 |
| Replicate B | -0.5166 | 0.3 |
| Replicate C | -1.1688 | 0.02 |

**Adjusted R<sup>2</sup>: 0.2272, p-value: $< 2.2 \times 10^{-16}$**
**Studentized Breusch-Pagan test p-value = 0.49**

**Extended Data Table 3.** Odds ratios (ORs) for the likelihood of identifying driver genes among mitosis-associated versus non-mitosis-associated genes in different tumor types. In each cell, the OR value is indicated, while the corresponding P values from Fisher’s exact tests are presented in parentheses. Asterisk (*) indicates statistical significance with a one-sided P value less than 0.05.

|  |  |  |  |  |  |  |  |  |  |  |  |
| --- | --- | --- | --- | --- | --- | --- | --- | --- | --- | --- | --- |
| All | 1.39<br>(0.127) | 1.19<br>(0.388) | 1.45<br>(0.138) | 1.68<br>(0.093) | 2.1<br>(0.002*) | 2.65<br>(0.035*) | 1.88<br>(0.038*) | 1.49<br>(0.131) | 2.47<br>(0.007*) | 0.41<br>(0.975) | 3.42<br>(9x<sup>-8</sup>*) |
| Mutations | 1.83<br>(0.045*) | 1.54<br>(0.214) | 1.65<br>(0.079) | 3.92<br>(0.026 *) | 2.68<br>(1x<sup>-4</sup> *) | 2.73<br>(0.046*) | 2.97<br>(0.006*) | 2.18<br>(0.041*) | 2.53<br>(0.011*) | 0.56<br>(0.832) | 2.5<br>(6x<sup>-4</sup>*) |
| mRNA | 1.88<br>(0.024*) | 1.36<br>(0.299) | 1.44<br>(0.154) | 3.88<br>(0.001 *) | 2.44<br>(5x<sup>-4</sup> *) | 2.41<br>(0.069*) | 2.9<br>(0.005*) | 1.96<br>(0.067) | 2.4<br>(0.014*) | 1.16<br>(0.489) | 3.61<br>(3x<sup>-8</sup>*) |
|  | BLCA | BRCA | COAD | ESCA | HNSC | KIRC | KIRP | LIHC | LUAD | PRAD | UCEC |

**Extended Data Table 4.** Primary Driver genes are evenly distributed across functional protein clusters. The odds ratios (ORs) indicate the likelihood of identifying driver genes within a specific functional cluster compared to others. Both two-sided p-values from Fisher’s exact tests and FDR-corrected p-values are provided. The analysis was conducted separately for all driver genes, and for two subsets: one with specific mutations, and another displaying altered expression in tumors.

| Tumor type | Cluster | All genes |  |  | Mut. genes |  |  | Expr. genes |  |  |
| --- | --- | --- | --- | --- | --- | --- | --- | --- | --- | --- |
|  |  | OR vs. other | Fisher exact test P | FDR | OR vs. other | Fisher exact test P | FDR | OR vs. other | Fisher exact test P | FDR |
| **BLCA** | mitotic cell cycle process | 1.771 | 0.297 | 1.000 | 1.048 | 1.000 | 1.000 | 1.784 | 0.402 | 0.917 |
| **BLCA** | rRNA metabolic process | 0.648 | 0.737 | 1.000 | 1.021 | 1.000 | 1.000 | 0.763 | 1.000 | 1.000 |
| **BLCA** | biological adhesion | 2.313 | 0.287 | 1.000 | 3.609 | 0.164 | 0.973 | 2.714 | 0.237 | 0.917 |
| **BLCA** | cation transport | 4.461 | 0.072 | 0.692 | 3.609 | 0.164 | 0.973 | 2.714 | 0.237 | 0.917 |
| **BLCA** | RNA splicing | 0.000 | 0.593 | 1.000 | 0.000 | 1.000 | 1.000 | 0.000 | 1.000 | 1.000 |
| **BLCA** | double-strand break repair via nonhomologous end joining | 1.093 | 1.000 | 1.000 | 1.659 | 0.507 | 1.000 | 1.271 | 0.591 | 0.929 |
| **BLCA** | mRNA transport | 0.000 | 1.000 | 1.000 | 0.000 | 1.000 | 1.000 | 0.000 | 1.000 | 1.000 |
| **BRCA** | mitotic cell cycle process | 0.231 | 0.240 | 1.000 | 0.281 | 0.400 | 1.000 | 0.281 | 0.400 | 0.917 |
| **BRCA** | rRNA metabolic process | 0.000 | 0.598 | 1.000 | 0.000 | 0.588 | 1.000 | 0.000 | 0.588 | 0.929 |
| **BRCA** | biological adhesion | 2.683 | 0.371 | 1.000 | 3.238 | 0.327 | 1.000 | 3.238 | 0.327 | 0.917 |
| **BRCA** | cation transport | 16.395 | 0.005 | 0.130 | 9.282 | 0.045 | 0.699 | 9.282 | 0.045 | 0.583 |
| **BRCA** | RNA splicing | 3.149 | 0.332 | 1.000 | 3.798 | 0.291 | 1.000 | 3.798 | 0.291 | 0.917 |
| **BRCA** | double-strand break repair via nonhomologous end joining | 3.149 | 0.332 | 1.000 | 3.798 | 0.291 | 1.000 | 3.798 | 0.291 | 0.917 |
| **BRCA** | mRNA transport | 0.000 | 1.000 | 1.000 | 0.000 | 1.000 | 1.000 | 0.000 | 1.000 | 1.000 |
| **COAD** | mitotic cell cycle process | 0.407 | 0.238 | 1.000 | 0.458 | 0.358 | 1.000 | 0.458 | 0.358 | 0.917 |
| **COAD** | rRNA metabolic process | 0.398 | 0.693 | 1.000 | 0.439 | 0.689 | 1.000 | 0.439 | 0.689 | 1.000 |
| **COAD** | biological adhesion | 11.835 | 0.004 | 0.130 | 13.372 | 0.003 | 0.202 | 13.372 | 0.003 | 0.202 |
| **COAD** | cation transport | 1.283 | 0.586 | 1.000 | 1.411 | 0.556 | 1.000 | 1.411 | 0.556 | 0.929 |
| **COAD** | RNA splicing | 3.933 | 0.150 | 0.975 | 4.356 | 0.130 | 0.969 | 4.356 | 0.130 | 0.917 |
| **COAD** | double-strand break repair via nonhomologous end joining | 1.508 | 0.536 | 1.000 | 1.659 | 0.507 | 1.000 | 1.659 | 0.507 | 0.929 |
| **COAD** | mRNA transport | 0.000 | 1.000 | 1.000 | 0.000 | 1.000 | 1.000 | 0.000 | 1.000 | 1.000 |
| **ESCA** | mitotic cell cycle process | 2.332 | 0.314 | 1.000 | 1.475 | 1.000 | 1.000 | 3.145 | 0.157 | 0.917 |
| **ESCA** | rRNA metabolic process | 0.547 | 1.000 | 1.000 | 0.000 | 1.000 | 1.000 | 0.000 | 0.354 | 0.917 |
| **ESCA** | biological adhesion | 1.752 | 0.488 | 1.000 | 5.425 | 0.230 | 1.000 | 1.986 | 0.451 | 0.917 |
| **ESCA** | cation transport | 4.572 | 0.120 | 0.921 | 5.425 | 0.230 | 1.000 | 5.254 | 0.099 | 0.877 |
| **ESCA** | RNA splicing | 0.000 | 1.000 | 1.000 | 0.000 | 1.000 | 1.000 | 0.000 | 1.000 | 1.000 |
| **ESCA** | double-strand break repair via nonhomologous end joining | 0.000 | 1.000 | 1.000 | 0.000 | 1.000 | 1.000 | 0.000 | 1.000 | 1.000 |
| **ESCA** | mRNA transport | 0.000 | 1.000 | 1.000 | 0.000 | 1.000 | 1.000 | 0.000 | 1.000 | 1.000 |
| **HNSC** | mitotic cell cycle process | 0.542 | 0.253 | 1.000 | 0.492 | 0.233 | 1.000 | 0.492 | 0.233 | 0.917 |
| **HNSC** | rRNA metabolic process | 1.966 | 0.226 | 1.000 | 1.647 | 0.359 | 1.000 | 1.647 | 0.359 | 0.917 |
| **HNSC** | biological adhesion | 2.799 | 0.170 | 0.975 | 3.151 | 0.138 | 0.969 | 3.151 | 0.138 | 0.917 |
| **HNSC** | cation transport | 4.912 | 0.040 | 0.499 | 5.553 | 0.029 | 0.602 | 5.553 | 0.029 | 0.481 |
| **HNSC** | RNA splicing | 1.790 | 0.615 | 1.000 | 2.007 | 0.345 | 1.000 | 2.007 | 0.345 | 0.917 |
| **HNSC** | double-strand break repair via nonhomologous end joining | 0.712 | 1.000 | 1.000 | 0.795 | 1.000 | 1.000 | 0.795 | 1.000 | 1.000 |
| **HNSC** | mRNA transport | 0.000 | 0.593 | 1.000 | 0.000 | 0.589 | 1.000 | 0.000 | 0.589 | 0.929 |
| **KIRC** | mitotic cell cycle process | 1.485 | 0.686 | 1.000 | 2.250 | 0.395 | 1.000 | 2.250 | 0.395 | 0.917 |
| **KIRC** | rRNA metabolic process | 1.020 | 1.000 | 1.000 | 0.000 | 0.590 | 1.000 | 0.000 | 0.590 | 0.929 |
| **KIRC** | biological adhesion | 0.000 | 1.000 | 1.000 | 0.000 | 1.000 | 1.000 | 0.000 | 1.000 | 1.000 |
| **KIRC** | cation transport | 9.282 | 0.045 | 0.499 | 12.351 | 0.031 | 0.602 | 12.351 | 0.031 | 0.481 |
| **KIRC** | RNA splicing | 0.000 | 1.000 | 1.000 | 0.000 | 1.000 | 1.000 | 0.000 | 1.000 | 1.000 |
| **KIRC** | double-strand break repair via nonhomologous end joining | 0.000 | 1.000 | 1.000 | 0.000 | 1.000 | 1.000 | 0.000 | 1.000 | 1.000 |
| **KIRC** | mRNA transport | 0.000 | 1.000 | 1.000 | 0.000 | 1.000 | 1.000 | 0.000 | 1.000 | 1.000 |
| **KIRP** | mitotic cell cycle process | 1.048 | 1.000 | 1.000 | 1.905 | 0.484 | 1.000 | 1.506 | 0.526 | 0.929 |
| **KIRP** | rRNA metabolic process | 1.021 | 1.000 | 1.000 | 0.000 | 0.354 | 1.000 | 0.000 | 0.366 | 0.917 |
| **KIRP** | biological adhesion | 1.411 | 0.556 | 1.000 | 1.986 | 0.451 | 1.000 | 1.752 | 0.488 | 0.917 |
| **KIRP** | cation transport | 7.185 | 0.027 | 0.420 | 5.254 | 0.099 | 0.846 | 9.329 | 0.016 | 0.407 |
| **KIRP** | RNA splicing | 0.000 | 1.000 | 1.000 | 0.000 | 1.000 | 1.000 | 0.000 | 1.000 | 1.000 |
| **KIRP** | double-strand break repair via nonhomologous end joining | 1.659 | 0.507 | 1.000 | 2.333 | 0.407 | 1.000 | 2.059 | 0.442 | 0.917 |
| **KIRP** | mRNA transport | 0.000 | 1.000 | 1.000 | 0.000 | 1.000 | 1.000 | 0.000 | 1.000 | 1.000 |
| **LIHC** | mitotic cell cycle process | 0.523 | 0.523 | 1.000 | 0.469 | 0.471 | 1.000 | 0.469 | 0.471 | 0.917 |
| **LIHC** | rRNA metabolic process | 0.000 | 0.209 | 1.000 | 0.000 | 0.352 | 1.000 | 0.000 | 0.352 | 0.917 |
| **LIHC** | biological adhesion | 8.126 | 0.021 | 0.407 | 6.159 | 0.079 | 0.764 | 6.159 | 0.079 | 0.874 |
| **LIHC** | cation transport | 15.327 | 0.002 | 0.130 | 13.123 | 0.008 | 0.303 | 13.123 | 0.008 | 0.303 |
| **LIHC** | RNA splicing | 0.000 | 1.000 | 1.000 | 0.000 | 1.000 | 1.000 | 0.000 | 1.000 | 1.000 |
| **LIHC** | double-strand break repair via nonhomologous end joining | 0.000 | 1.000 | 1.000 | 0.000 | 1.000 | 1.000 | 0.000 | 1.000 | 1.000 |
| **LIHC** | mRNA transport | 2.221 | 0.423 | 1.000 | 3.238 | 0.327 | 1.000 | 3.238 | 0.327 | 0.917 |
| **LUAD** | mitotic cell cycle process | 0.710 | 0.761 | 1.000 | 0.606 | 0.739 | 1.000 | 0.606 | 0.739 | 1.000 |
| **LUAD** | rRNA metabolic process | 1.021 | 1.000 | 1.000 | 1.300 | 0.668 | 1.000 | 1.300 | 0.668 | 1.000 |
| **LUAD** | biological adhesion | 1.411 | 0.556 | 1.000 | 1.752 | 0.488 | 1.000 | 1.752 | 0.488 | 0.917 |
| **LUAD** | cation transport | 3.609 | 0.164 | 0.975 | 1.752 | 0.488 | 1.000 | 1.752 | 0.488 | 0.917 |
| **LUAD** | RNA splicing | 1.659 | 0.507 | 1.000 | 2.059 | 0.442 | 1.000 | 2.059 | 0.442 | 0.917 |
| **LUAD** | double-strand break repair via nonhomologous end joining | 1.659 | 0.507 | 1.000 | 2.059 | 0.442 | 1.000 | 2.059 | 0.442 | 0.917 |
| **LUAD** | mRNA transport | 2.004 | 0.453 | 1.000 | 2.486 | 0.392 | 1.000 | 2.486 | 0.392 | 0.917 |
| **PRAD** | mitotic cell cycle process | 0.727 | 1.000 | 1.000 | 0.000 | 1.000 | 1.000 | 0.727 | 1.000 | 1.000 |
| **PRAD** | rRNA metabolic process | 0.000 | 1.000 | 1.000 | 0.000 | 1.000 | 1.000 | 0.000 | 1.000 | 1.000 |
| **PRAD** | biological adhesion | 0.000 | 1.000 | 1.000 | 0.000 | 1.000 | 1.000 | 0.000 | 1.000 | 1.000 |
| **PRAD** | cation transport | 8.112 | 0.177 | 0.975 | Inf | 0.063 | 0.764 | 8.112 | 0.177 | 0.917 |
| **PRAD** | RNA splicing | 0.000 | 1.000 | 1.000 | 0.000 | 1.000 | 1.000 | 0.000 | 1.000 | 1.000 |
| **PRAD** | double-strand<br>break repair<br>via<br>nonhomologou<br>s end joining | 0.000 | 1.000 | 1.000 | 0.000 | 1.000 | 1.000 | 0.000 | 1.000 | 1.000 |
| **PRAD** | mRNA<br>transport | 0.000 | 1.000 | 1.000 | 0.000 | 1.000 | 1.000 | 0.000 | 1.000 | 1.000 |
| **UCEC** | mitotic cell<br>cycle process | 1.964 | 0.103 | 0.877 | 0.755 | 0.629 | 1.000 | 1.964 | 0.103 | 0.877 |
| **UCEC** | rRNA<br>metabolic<br>process | 0.666 | 0.591 | 1.000 | 0.883 | 1.000 | 1.000 | 0.666 | 0.591 | 0.929 |
| **UCEC** | biological<br>adhesion | 1.000 | 1.000 | 1.000 | 1.878 | 0.610 | 1.000 | 1.000 | 1.000 | 1.000 |
| **UCEC** | cation<br>transport | 1.872 | 0.412 | 1.000 | 1.878 | 0.610 | 1.000 | 1.872 | 0.412 | 0.917 |
| **UCEC** | RNA splicing | 1.211 | 1.000 | 1.000 | 2.270 | 0.300 | 1.000 | 1.211 | 1.000 | 1.000 |
| **UCEC** | double-strand<br>break repair<br>via<br>nonhomologou<br>s end joining | 2.360 | 0.365 | 1.000 | 4.508 | 0.076 | 0.764 | 2.360 | 0.365 | 0.917 |
| **UCEC** | mRNA<br>transport | 0.000 | 0.336 | 1.000 | 0.000 | 0.589 | 1.000 | 0.000 | 0.336 | 0.917 |

**Extended Data Table 5.** Primary antibodies used for the immunofluorescence stainings.

| **Antibody HPA ID** | **Protein** | **Species** | **Concentration (mg/mL)** | **Dilution** |
| --- | --- | --- | --- | --- |
| HPA004748 | TLN1 | Rabbit | 0.06 | 1:30 |
| HPA030052 | EPRS1 | Rabbit | 0.0963 | 1:50 |
| HPA032151 | DIAPH3 | Rabbit | 0.2773 | 1:140 |
| HPA034506 | MAPK1IP1L | Rabbit | 0.0793 | 1:40 |
| HPA034569 | UBE2C | Rabbit | 0.5514 | 1:200 |
| HPA041354 | FLYWCH2 | Rabbit | 0.1076 | 1:55 |
| HPA042123 | SUMO2 | Rabbit | 0.1632 | 1:80 |
| HPA042904 | NUSAP1 | Rabbit | 0.7772 | 1:200 |
| HPA045034 | PTTG1 | Rabbit | 0.2154 | 1:100 |
| HPA047929 | PCLAF | Rabbit | 0.0264 | 1:7 |
| HPA052206 | TFG | Rabbit | 0.1796 | 1:90 |
| HPA054319 | GPATCH4 | Rabbit | 0.2894 | 1:150 |
| HPA061291 | C19orf53 | Rabbit | 0.1424 | 1:70 |
| HPA061448 | CCNB1 | Rabbit | 0.792 | 1:200 |
| HPA070550 | INCENP | Rabbit | 0.3649 | 1:150 |

